# Goals shape human confidence

**DOI:** 10.64898/2026.09.02.748915

**Authors:** Pradyumna Sepulveda, Stephen M. Fleming, Mariana Zurita, Benedetto de Martino

**Affiliations:** Department of Psychiatry, Vagelos College of Physicians and Surgeons, Columbia University, New York, NY, USA; Institute of Cognitive Neuroscience, University College London, London, United Kingdom; Max Planck UCL Centre for Computational Psychiatry and Aging Research, University College London, United Kingdom; Department of Experimental Psychology, University College London, United Kingdom

## Abstract

Confidence guides our decisions: it determines when we commit to a choice, gather more information, or change our minds. Prominent models treat confidence as a readout of the evidence in support of a decision: given the same evidence, a similar confidence level should follow, regardless of what the decision-maker is trying to achieve. Here we show instead that confidence formation is relative to the goals of the decision-maker. Across value-based and perceptual decisions in different sensory modalities, we manipulated the behavioural goal, requiring participants to select the option with either the most or the least evidence. Reversing the goal reversed the relationship between evidence and confidence: biases previously considered fixed features of a confidence computation, such as the positive evidence bias (PEB), followed the goal rather than the stimulus. We developed a new computational model, GOAL (Goal-Oriented Asymmetric Likelihood), in which evidence is weighted according to its relevance for the current goal, and validated it in behavioural and physiological (pupillometry) experiments. Our findings support a new account of metacognition in which confidence is constructed relative to the goals of the decision-maker, prioritising goal-supportive evidence.

## 1 Introduction

Decisions about perceptual features or personal preferences are often accompanied by an internal sense of confidence. Confidence plays a fundamental role in allowing us to assess past actions (e.g., “I’m sure I did not miss the red light!”) and to inform future choices (e.g., “I’m quite confident I’ll come back to this restaurant”). In both cases confidence is useful precisely because it serves 1 the goals of the agent: it tells us whether we are likely to have achieved what we set out to do. Yet the goals of the decision-maker are absent from most formal accounts of how confidence is computed.

According to standard models, confidence reflects the probability that a decision is correct,^1^ typically determined by the balance of evidence supporting or opposing that decision.^2,3^ In this framework the mapping from evidence to confidence is fixed: given the same stimulus and the same choice, an observer should report a similar level of confidence regardless of what they are trying to achieve. Confidence, in other words, is blind to the goal of the decision-maker.

Recent research has shown puzzling departures from these predictions.^4–7^ One well-documented deviation is the positive evidence bias (PEB): a tendency for confidence to overemphasise evidence favouring the chosen option while underweighting evidence for the unchosen one.^8–13^ A related phenomenon is that confidence increases with the sum of evidence across all available options (ΣEvidence), rather than only with the balance between them.^14–17^ Computational models developed to explain the PEB also predict this ΣEvidence effect, linking both to a detection-like process in which confidence is driven by stimulus features such as the overall strength or visibility of the perceptual evidence.^10,18–20^

These biases are usually treated as quirks of a confidence computation: fixed distortions rooted in the perceptual properties of the stimulus. Critically, however, in most experiments in which these biases have been observed, the observer has a single goal: selecting the option supported by stronger evidence. An alternative possibility, then, is that the PEB and related biases are due to the observer’s goal shaping how evidence is used to inform confidence. Unless the goal is explicitly manipulated, it is not possible to disentangle whether biases in confidence are driven by the stimulus perception or the goal. Our recent work shows that goals and context strongly influence how people sample and accumulate information,^15^ raising the possibility that the same is true of how confidence is formed. If so, phenomena such as the PEB and the ΣEvidence effect would not be fixed features of metacognition but signatures of a more general, goal-relative computation. Testing between these two accounts is the main aim of the present study.

To do so, we designed a series of experiments that decoupled the decision-maker’s goal from the evidence supporting a decision, across both perceptual (in different sensory modalities) and value-based domains. Alongside standard “high-evidence” decisions, in which participants chose the option with more evidence, we introduced “low-evidence” decisions, requiring participants to select the option with less evidence. The two accounts make opposite predictions: if confidence is a fixed function of the stimulus, the relationship between evidence and confidence should be unchanged by the goal; if confidence is goal-relative, this relationship should be reversed. We found the latter: in high-evidence tasks, confidence increased with both chosen evidence and the overall evidence available in the choice (as previously reported), whereas in low-evidence tasks these relationships inverted, with confidence decreasing as evidence strength increased. Thus, under low-evidence goals, weaker evidence was associated with greater confidence, ruling out an account driven solely by the perceptual properties of the stimuli.

To track the formation of these goal-dependent confidence signals, we complemented behavioural data with physiological measures of pupil dilation. Variation in pupil size has been employed extensively to study the unfolding of cognitive processes over time.^21,22^ The dilation of pupils is controlled by the noradrenergic locus coeruleus (LC), a small brainstem nucleus with an important role in task-related processes,^23^ and increases in response to effort, low-confidence choices, and higher levels of uncertainty and surprise.^24–30^ Pupil-linked arousal thus offers an independent window onto confidence formation, revealing whether goal-dependent shifts in reported confidence track corresponding shifts in decision-related arousal.

The same reversal emerged across perceptual and value-based domains and was accompanied by matching changes in pupil size, consistent with a goal-dependent construction of confidence. To capture the computational roots of these effects, we developed a new model grounded in signal detection theory,^31^ which we call GOAL (Goal-Oriented Asymmetric Likelihood). Based on an asymmetric evaluation of evidence according to its relevance to the current goal, the model captures the reversal of the ΣEvidence effect and recasts the PEB as a special case, one that emerges whenever the goal happens to be selecting the stronger option. Taken together, our results indicate that goal-dependent effects on confidence are not anomalies but a fundamental feature of human metacognition, in which confidence is dynamically constructed to align with the goals of the decision-maker.

## 2 Results

We conducted a series of decision-making experiments to investigate how goals influence the construction of confidence. We reanalysed two previously published datasets^15^ and combined them with results from two newly conducted experiments. Across all four experiments, participants made either value-based or perceptual decisions within one of two task frames: in a positive frame, they selected the option with higher evidence, and in a negative frame, they selected the option with lower evidence. After each decision, participants reported their level of confidence in their choice.

### 2.1 Effects of evidence biases on confidence depend on frame

#### 2.1.1 Experiment 1

In the first value-based decision experiment (*n* = 31), hungry participants chose between pairs of snack items under two framing conditions: a *like* frame, selecting the snack they preferred to eat, and a *dislike* frame, the one they wished to avoid (Figure 1A). After each choice, they rated their confidence, and before the task provided subjective value ratings.^32^ Using this design we tested how task goals influence confidence in value-based choices. To test for a positive evidence bias (PEB) we fitted a hierarchical linear model predicting confidence from the evidence supporting the chosen and unchosen options (i.e., participants’ subjective values for each snack). In the *like* frame (blue), confidence increased with higher chosen evidence (*β*_chosen_ = 0.21 ± 0.03, *p <* 0.001) and decreased with higher unchosen evidence (*β*_unchosen_ = −0.07 ± 0.03, *p <* 0.01) (Figure 1B). The effect of chosen evidence was significantly stronger than that of unchosen evidence (|*β*_chosen_| = 0.22 vs |*β*_unchosen_| = 0.10; *t*(30) = 5.002, *p <* 0.001), confirming an overweighting of the selected alternative, and consistent with previous reports of a PEB.^8–13,17^ Reaction time also negatively predicted confidence (*β*_RT_ = −0.26 ± 0.03, *p <* 0.001).

**Figure 1:**
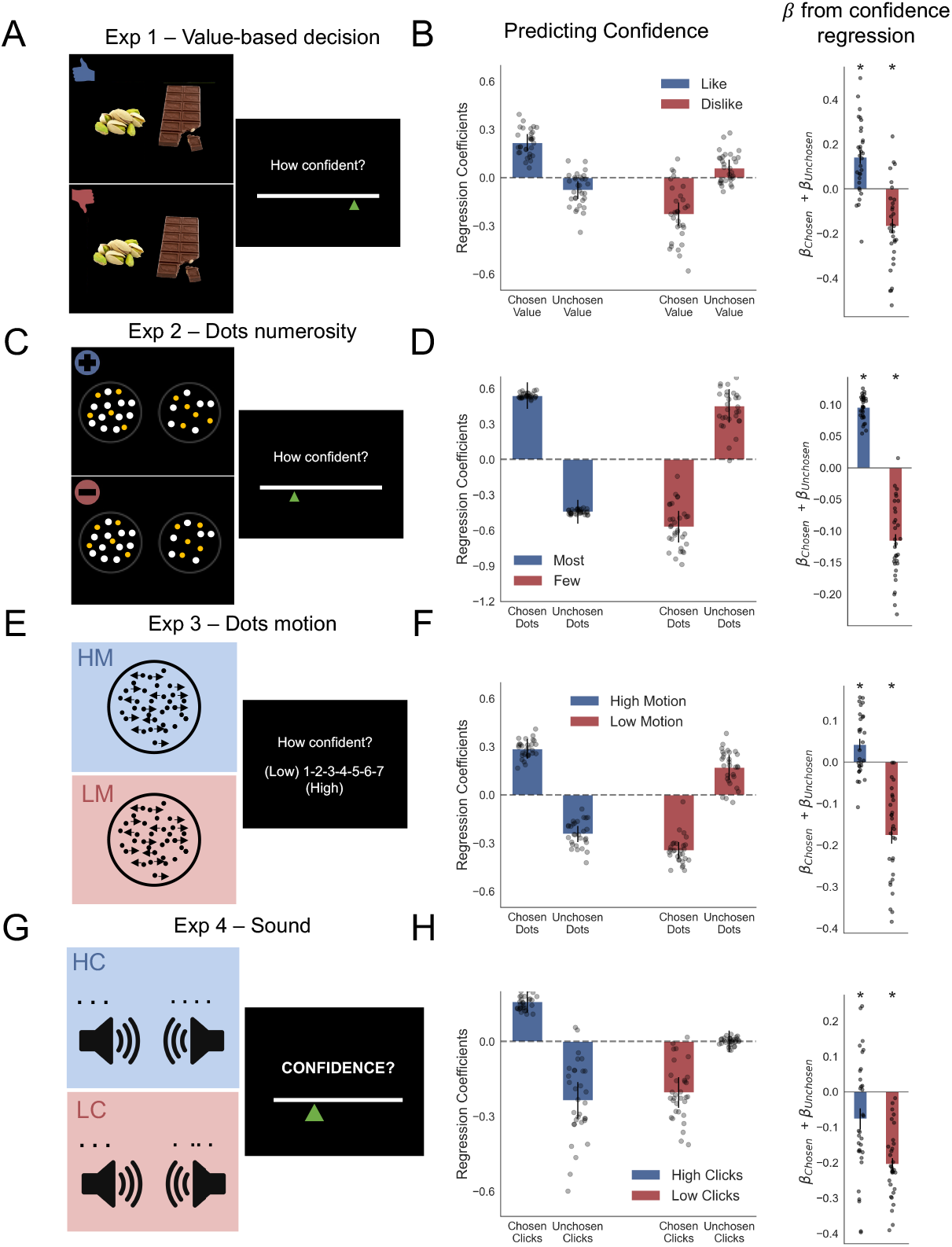
Goal-dependent asymmetries in evidence integration for value-based and perceptual decision confidence. (A) Experiment 1 – Value-based decisions. Hungry participants chose between pairs of snack items, either selecting the item they preferred to eat (*like* frame) or the one they wished to avoid (*dislike* frame). Subjective values for each item were recorded before the decision task using the Becker–DeGroot–Marschak procedure. (B) Regression results of Experiment 1. Confidence depended asymmetrically on chosen and unchosen evidence, and this asymmetry reversed across frames. In the *like* frame, higher chosen value increased confidence, whereas in the *dislike* frame it decreased confidence. The overall evidence effect (*β*_chosen_ +*β*_unchosen_) was positive in the *like* frame and negative in the *dislike* frame, reflecting a goal-dependent reversal. (C) Experiment 2 – Dot numerosity. Participants selected between two circles containing different numbers of dots. In the *most* frame they chose the circle with more dots, and in the *fewest* frame the one with fewer dots. (D) Regression results of Experiment 2. Confidence again showed an evidence asymmetry modulated by task frame. Chosen evidence had a stronger influence on confidence than unchosen evidence, with a positive overall effect in the *most* frame and a negative one in the *fewest* frame. (E) Experiment 3 – Dot motion. Participants judged the dominant motion direction (left or right) in a random dot kinematogram. In the *High Motion* frame, they selected the direction with more coherently moving dots; in the *Low Motion* frame, the direction with fewer. (F) Regression results of Experiment 3. Confidence patterns mirrored those of the previous experiments: in the *High Motion* frame, chosen evidence boosted confidence, while in the *Low Motion* frame it reduced it. (G) Experiment 4 – Auditory clicks. Participants listened to stereo click sequences and indicated the side with more (*High Clicks*) or fewer (*Low Clicks*) clicks. (H) Regression results of Experiment 4. Confidence in the *Low Clicks* frame followed the same pattern as in earlier tasks, while in the *High Clicks* frame the unchosen evidence contributed slightly more. Nonetheless, the overall evidence–confidence relationship remained frame-dependent, with a stronger negative effect in the *Low Clicks* frame. Across all experiments, confidence was reported after each decision. In regression plots, bars represent fixed effects and dots represent mixed effects; error bars denote 95% confidence intervals for fixed effects. Significance levels: *p <* 0.05; *p <* 0.01; *p <* 0.001.

Our framing manipulation allowed us to decouple positive evidence (in this case positive value evidence) from choices, since in the *dislike* frame participants were instructed to choose the snack they least preferred. We showed that, contrary to the pattern observed in the *like* frame, confidence decreased with chosen evidence (*β*_chosen_ = −0.22 ± 0.03, *p <* 0.001) and increased with unchosen evidence (*β*_unchosen_ = 0.06 ± 0.03, *p <* 0.05). Again, the chosen option dominated confidence judgments (|*β*_chosen_| = 0.24 vs |*β*_unchosen_| = 0.09; *t*(30) = 5.783, *p <* 0.001). As in the *like* frame, reaction time negatively predicted confidence (*β*_RT_ = −0.29 ± 0.04, *p <* 0.001).

These results indicate that PEB is not simply driven by the presence of positive evidence, but by how evidence is used relative to the goal of the task. This goal-dependent modulation becomes clearer when combining the chosen and unchosen coefficients to estimate the overall evidence effect (*β*_overall_ = *β*_chosen_ + *β*_unchosen_). The overall evidence had opposite influences on confidence in the two frames (*like*: *β*_overall_ = 0.14 ± 0.03; *dislike*: *β*_overall_ = −0.17 ± 0.03; *t*(30) = 8.961, *p <* 0.001) (Figure 1B). In the *dislike* frame, greater evidence for the chosen option actually reduced confidence, revealing a negative evidence bias.

#### 2.1.2 Experiment 2

In the second, perceptual discrimination experiment (*n* = 32), participants viewed two circles containing different numbers of dots (Figure 1C). In the *most* frame, they identified the circle with more dots, and in the *fewest* frame, the circle with fewer dots. We observed a comparable behavioural pattern to Experiment 1, which extended the findings to the perceptual domain. Participants’ confidence systematically overweighted the evidence favouring the chosen option (number of dots) relative to the unchosen option in both the *most* and *fewest* frames (most: |*β*_chosen_| = 0.54, |*β*_unchosen_| = 0.44, *t*(31) = 28.78, *p <* 0.001; fewest: |*β*_chosen_| = 0.57, |*β*_unchosen_| = 0.45, *t*(31) = 10.887, *p <* 0.001).

Critically, and similar to the pattern observed in Experiment 1, the direction of the effect was modulated by the frame. In the *most* frame, higher evidence for the chosen patch increased confidence (*β*_chosen_ = 0.53 ± 0.06, *p <* 0.001), whereas in the *fewest* frame, the same increase in chosen evidence reduced confidence (*β*_chosen_ = −0.56 ± 0.07, *p <* 0.001). The opposite pattern was observed for the unchosen patch (*most*: *β*_unchosen_ = −0.44 ± 0.05, *p <* 0.001; *fewest*: *β*_unchosen_ = 0.45 ± 0.07, *p <* 0.001) (Figure 1D).

These results demonstrate that a positive evidence bias in perceptual confidence is also shaped by the decision goal, mirroring the findings from the value-based task. This goal dependence was further confirmed by the reversal of the overall evidence effect on confidence between frames (*most*: *β*_overall_ = 0.10 ± 0.003; *fewest*: *β*_overall_ = −0.12 ± 0.01, *t*(31) = 18.341, *p <* 0.001). In the *fewest* frame, confidence increased when less physical evidence (fewer dots) was presented.

As in Experiment 1, the framing manipulation did not alter the sign of the reaction time effect: confidence consistently decreased with longer RTs (*most*: *β*_RT_ = −0.32 ± 0.03, *p <* 0.001; *fewest*: *β*_RT_ = −0.27 ± 0.03, *p <* 0.001).

#### 2.1.3 Experiment 3

The goal of Experiment 3 was to test whether the effects observed in Experiment 1 and 2 were still present, even when controlling for shifts in the spatial allocation of attention (i.e., gaze shifts between left and right stimuli). To this end, we used a dot motion task, which has previously been shown to elicit a positive evidence bias (PEB)^8,9,16^ (Figure 1E). The motion discrimination task, which presented all evidence within a single central aperture, allowed us to test if the effect we observed was also present when supporting and opposing evidence appeared at the same spatial location. While in the classic random dot kinematogram (RDK), a subset of dots moves coherently in a single target direction while the remaining “noise” dots move independently in random directions (typically drawn from the full 360*^◦^* range), in our RDK we constrained dot motion to only rightward or leftward directions (0*^◦^* and 180*^◦^*directions, respectively). This was done to clearly measure evidence in favour of and against participants’ choices. Participants were asked to judge the dominant motion direction (left or right) in a random dot kinematogram. In the *high motion* frame, they were asked to report the direction with more coherently moving dots; in the *low motion* frame, the direction with fewer moving dots.

As in the previous experiments, we fitted a hierarchical linear model predicting confidence from the evidence supporting the chosen and unchosen directions (Figure 1F). Once again, confidence overweighted the evidence for the chosen relative to the unchosen direction in both the *high motion* (|*β*_chosen_| = 0.29, |*β*_unchosen_| = 0.24, *t*(28) = 3.135, *p <* 0.01) and *low motion* frames (|*β*_chosen_| = 0.35, |*β*_unchosen_| = 0.18, *t*(28) = 8.787, *p <* 0.001).

As in the previous experiments, the direction of the chosen-evidence effect reversed across frames: confidence increased with greater motion evidence in the *high motion* frame (*β*_chosen_ = 0.28 ± 0.03, *p <* 0.001) but decreased with motion evidence in the *low motion* frame (*β*_chosen_ = −0.36 ± 0.03, *p <* 0.001). The opposite pattern was observed for unchosen evidence (*high motion*: *β*_unchosen_ = −0.24 ± 0.025, *p <* 0.001; *low motion*: *β*_unchosen_ = 0.17 ± 0.04, *p <* 0.001). In other words, while confidence was enhanced when more dots moved in the chosen direction during the *high motion* frame, it increased when fewer dots moved during the *low motion* frame.

The overall evidence effect on confidence also flipped across frames (*high motion*: *β*_overall_ = 0.04 ± 0.013; *low motion*: *β*_overall_ = −0.18 ± 0.02; *t*(28) = 9.625, *p <* 0.001), confirming a strong goal-dependent modulation. Reaction time effects were again consistent across frames, with longer RTs predicting lower confidence (*high motion*: *β*_RT_ = −0.31 ± 0.03, *p <* 0.001; *low motion*: *β*_RT_ = −0.29 ± 0.024, *p <* 0.001).

These findings demonstrate that a goal-dependent evidence bias in confidence is robustly elicited even when controlling for shifts in spatial attention.

#### 2.1.4 Experiment 4

The fourth experiment extended our investigation to the auditory domain while recording participants’ pupil diameter (*n* = 32), as an index of arousal-related processes previously linked to decision confidence and uncertainty.^25,26^ Participants viewed a constant-luminance screen while hearing click sequences played simultaneously to the left and right ears (Figure 1G). In the *high clicks* frame, they selected the side with more clicks, and in the *low clicks* frame, the side with fewer clicks. This design allowed us to test whether goal-dependent confidence effects generalise beyond visual tasks, while pupillometry let us examine whether these effects are reflected in physiological signatures of the decision process (see below for details on pupil analysis).

We fitted a hierarchical linear model predicting confidence from the evidence supporting the chosen and unchosen options, defined as the number of clicks presented in the left and right speakers (Figure 1H). In the *low clicks* frame, we observed a strong asymmetry in evidence integration: confidence was primarily influenced by the chosen evidence (|*β*_chosen_| = 0.21; |*β*_unchosen_| = 0.01; *t*(31) = 9.987, *p <* 0.001). In contrast, in the *high clicks* frame, this asymmetry reversed: unchosen evidence had a stronger effect than chosen evidence (|*β*_chosen_| = 0.16; |*β*_unchosen_| = 0.24; *t*(31) = −2.981, *p <* 0.001).

Despite these differences, the direction of the chosen-evidence effect remained consistent with previous experiments: confidence increased with more clicks on the chosen side in the *high clicks* frame and decreased in the *low clicks* frame (*high clicks*: *β*_chosen_ = 0.15 ± 0.02, *p <* 0.001; *low clicks*: *β*_chosen_ = −0.20 ± 0.03, *p <* 0.001). Unchosen evidence negatively affected confidence in the *high clicks* frame (*β*_unchosen_ = −0.23 ± 0.04, *p <* 0.001) but showed no reliable effect in the *low clicks* frame (*β*_unchosen_ = 0.00 ± 0.02, *p* = 0.99, n.s.).

The overall evidence effect on confidence differed significantly between frames, with a stronger negative relationship in the *low clicks* frame (*high clicks*: *β*_overall_ = −0.08 ± 0.03; *low clicks*: *β*_overall_ = −0.20 ± 0.02; *t*(32) = 5.851, *p <* 0.001). Thus, as in the visual tasks, the decision frame modulated how evidence shaped confidence, though here the influence of low evidence (i.e., fewer auditory events) was more pronounced in the negative frame.

We attribute the absence of a positive overall chosen-evidence effect (*β*_chosen_ + *β*_unchosen_) in the *high clicks* frame to differences in task structure in the auditory Experiment 4. Unlike the visual Experiments 2 and 3, where participants could freely sample visual evidence until making a choice, auditory evidence was restricted to a 1-second presentation window. Although participants were allowed unlimited time to respond, no further evidence was available. Previous research has shown that such temporal constraints can weaken the relationship between confidence and other behavioural measures, such as reaction time,^4^ which may also explain the reduced positive evidence bias observed here.

As in all previous experiments, the framing manipulation did not alter the direction of the reaction time effect: confidence consistently decreased with longer RTs (*high clicks*: *β*_RT_ = −0.29 ± 0.03, *p <* 0.001; *low clicks*: *β*_RT_ = −0.34 ± 0.03, *p <* 0.001).

Overall, despite the absence of a clear positive evidence bias in the *high clicks* frame, the results of Experiment 4 further confirm that the goal framing modulates the relationship between evidence and confidence, with confidence in the low clicks frame being driven more strongly by the absence of evidence (i.e., fewer auditory events).

### 2.2 Asymmetric variance generates a goal-dependent evidence bias

The previous analyses revealed a robust, domain-general goal-based evidence bias in confidence. To uncover the computational mechanisms underlying this effect, we developed a Bayesian graphical model grounded in signal detection theory (SDT) in which we assume that the observer compares the sample they observe to two internal belief distributions: one for high evidence and another for low evidence (Figure 2A). The core assumption is that the observer compares the available evidence on each trial with their internal belief distributions to select the combination that is most probable. For example, deciding whether the left option belongs to the low-evidence distribution and the right to the high-evidence one, or vice versa.

**Figure 2:**
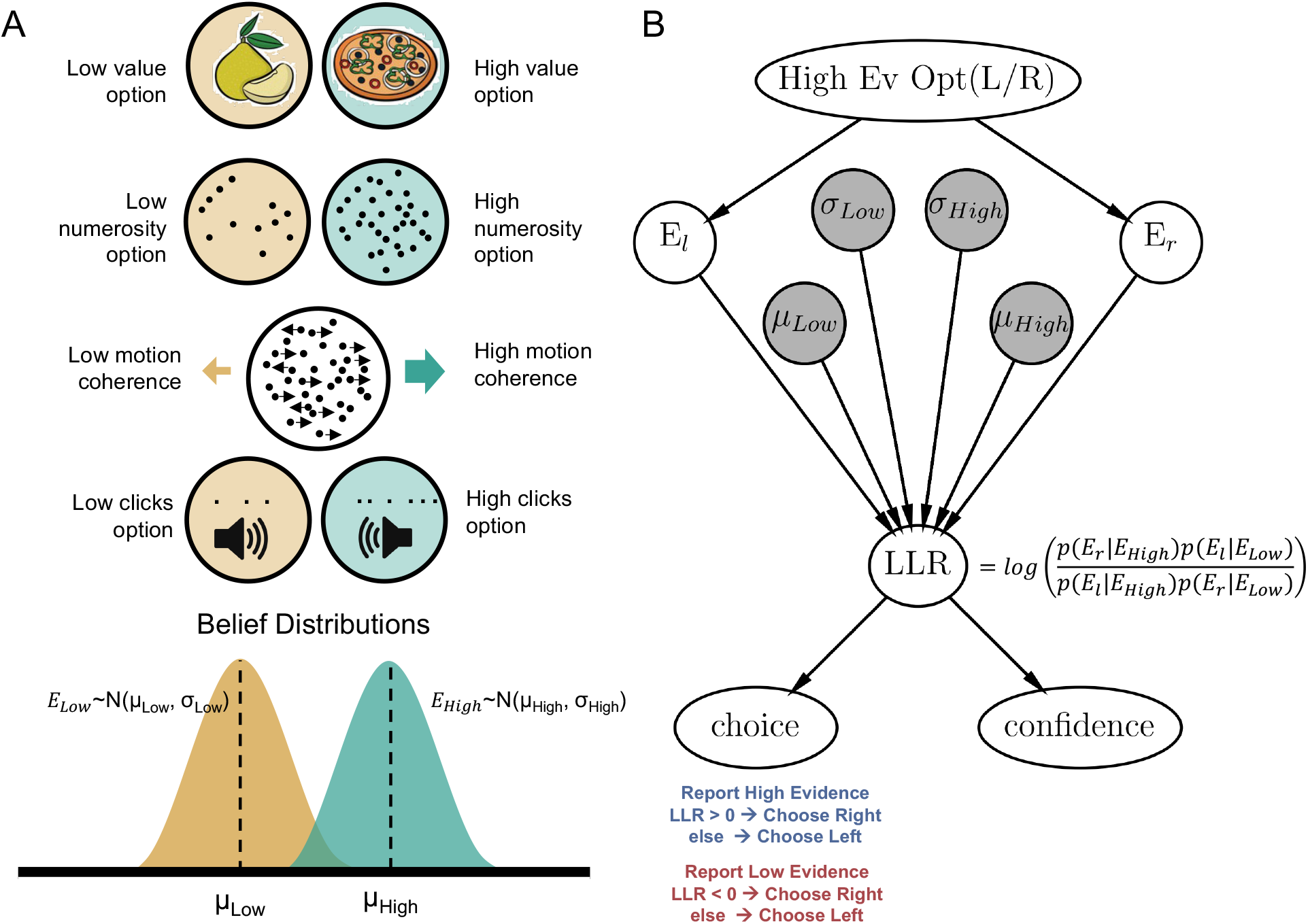
Bayesian observer model – Goal-oriented Asymmetric Likelihood (GOAL). (A) The model proposed in this work is grounded in SDT. GOAL considers an observer with internal belief distributions that categorise the alternatives as high evidence and low evidence options. These two belief distributions were modelled as Gaussians with parameters (*µ*_high_*, σ*_high_) and (*µ*_low_*, σ*_low_). This separation into two categories can be applied to the evidence for each of our experiments. As stated in its name, GOAL allows variances to vary independently, which captures the evidence effects on confidence observed in human experiments and its task-goal dependency. In high evidence frames (e.g., *like, most, high motion*, or *high clicks* frames) the model predicts *σ*_high_ *> σ*_low_. In low evidence frames, *σ*_high_ *< σ*_low_ can generate the behavioural effects found in humans. (B) Bayesian graphical model used for simulations. For details, see Methods – Model simulations. High Ev Opt (L/R): for simulations, the direction of high evidence was generated from random samples of a Bernoulli distribution; *E_r_*: observed evidence for the right-side alternative; *E_l_*: observed evidence for the left-side alternative; LLR: log-likelihood ratio term (*L*).

This comparison is formalised as a decision variable, expressed as a log-likelihood ratio (*L*). Assuming the goal is to report the option with higher evidence, when *L* ≥ 0, the observer selects the right option; and when *L <* 0, they select the left option. Crucially, the mapping between log-likelihood ratio and choice is goal-dependent: in low-evidence frames, this rule reverses, with *L* ≥ 0 leading to a choice of the left option, and *L <* 0 the right (Figure 2B).

For the following section we present SDT-inspired models that try to capture choice and confidence behaviour. For each one of the following models, we simulated 2,000 trials and fitted logistic and linear regressions to synthetic choice and confidence data, respectively, across both high– and low-evidence frames. Full details of the simulation procedures are provided in the Methods section.

#### 2.2.1 Standard equal variance model does not capture biases in confidence

The classical approach for the modelling of decision confidence is SDT, which assumes equal variance for the target and non-target evidence distributions.^10,31,33–35^ In our experiments, we call this instantiation the Equal Variance Model (EVM), where we consider variance values for high and low evidence distributions to be identical. EVM simulations reproduced the basic frame-dependent choice pattern (Figure 3A, top). The step-like shape of the logistic curve arose from the deterministic decision threshold (i.e., when *L* ≥ 0 the right option was chosen, otherwise the left). Confidence was computed as the magnitude of the likelihood ratio (|*L*|), following the same rule for both simulated frames. Simulated confidence showed the expected U-shape as a function of the evidence difference between options, which is a typical signature of human confidence reports^36^ (Figure 3A, bottom). However, the model’s patterns of confidence diverged sharply from the human data: a linear regression predicting confidence showed that chosen and unchosen evidence contributed with equal and opposite weight in both frames (high frame: *β*_chosen_ = 0.846, *p <* .001, *β*_unchosen_ = −0.859, *p <* .001; low frame: *β*_chosen_ = −0.854, *p <* .001, *β*_unchosen_ = 0.856, *p <* .001; Figure 3B). Because these weights were symmetric, the model predicted no frame modulation of the overall evidence effect (*β*_chosen_ + *β*_unchosen_) on confidence (high frame: *β*_overall_ = 4.83 × 10*^−^*^5^, *p* = .980, ns; low frame: *β*_overall_ = 57.31 × 10*^−^*^5^, *p* = .75, ns; Figure 3C).

**Figure 3:**
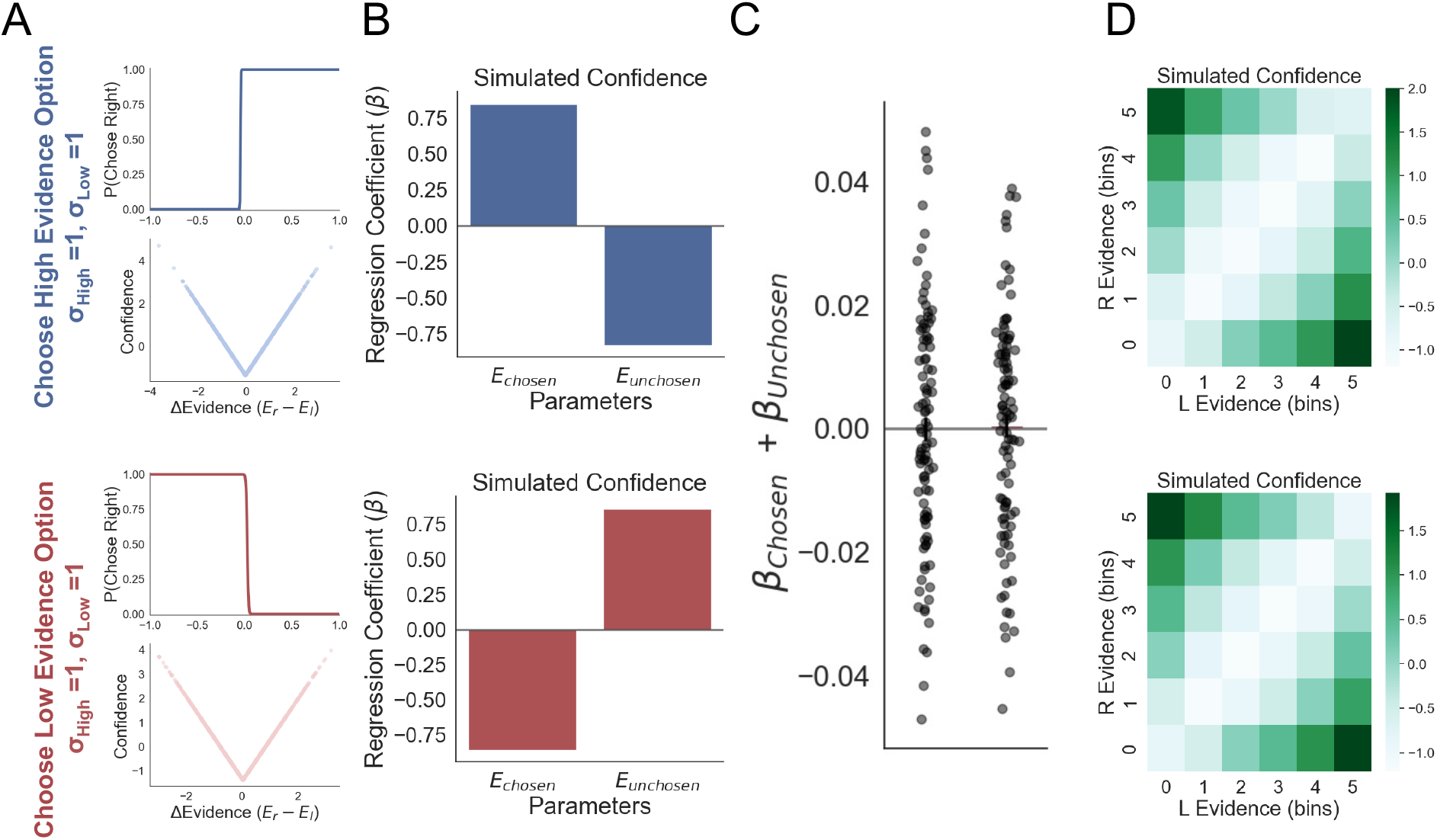
Equal Variance Model (EVM) simulations. Frames in which the model selects high evidence (blue) or low evidence (red) are presented. (A) Simulated choice and confidence. (B) Linear regression coefficients of variables predicting confidence using chosen and unchosen evidence as predictors indicated an equal weight of both streams of evidence. (C) The overall evidence effect (*β*_chosen_ + *β*_unchosen_) from simulated trials did not show an asymmetry in the evidence integration between high and low evidence frames. (D) Confidence in the evidence space, defined by (binned) evidence for left and right options.

For visualisation, we plotted confidence across the evidence space, defined by the evidence strength of the left and right options (Figure 3D). Lower confidence values appeared over the diagonal of the evidence-space. The evidence-space plot showed no frame-dependent displacement of confidence for the EVM simulated confidence reports (Figure 3D). This pattern held across a range of equal-variance values (*σ*_high_ = *σ*_low_; Supplemental Figure S1), confirming it is a structural property of the equal-variance assumption rather than a specific parameter choice. In other words, the EVM behaves like a confidence signal generated by a simple balance-of-evidence rule.^10,35^ This is insufficient to capture the behavioural results in which participants’ confidence showed a goal-dependent overall evidence effect. This discrepancy indicates that an additional mechanism is required to explain how confidence is shaped by task goals.

#### 2.2.2 Goal-oriented Asymmetric Likelihood Model (GOAL)

We reasoned that this missing mechanism might lie in how the precision of belief, rather than only its mean, changes with the task goal. Previous work suggests that unequal variances between target and non-target distributions, specifically a higher variance for the target distribution 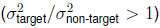, provide a more realistic description of decision-making in ecological contexts, or in tasks where the target signal must be distinguished from a noisy background.^37,38^ This asymmetry has a plausible origin in the way neural response variability scales with signal strength:^39^ when a stimulus carries more of the target signal (e.g., more visual dots, higher luminosity) the internal representation of that information tends to be proportionally noisier, consistent with Poisson-like scaling of neuronal firing, where response variance increases with response magnitude.^40,41^

Building on this precedent, we reasoned that the same logic could extend naturally to our goal-manipulation. In classical unequal-variance SDT, the identity of the higher-variance distribution is fixed, because it always corresponds to the physical signal (e.g., “target present”). In our task, however, what counts as the informative, the goal-relevant category, is not fixed by the stimulus but is set by instruction: participants must extract graded information about the high-evidence option when asked to report high evidence, and about the low-evidence option when asked to report low evidence. If higher variance reflects the amount of information an observer must extract from a category, rather than a fixed physical property of the stimulus, then whichever category is currently goal-relevant should inherit the “higher-variance status.” We therefore built the Goal-Oriented Asymmetric Likelihood (GOAL), a model similar to standard SDT but with asymmetric variance that depends on the task goal. Therefore, in the GOAL model the decision-maker changes the relative variance of the internal belief distributions against which they compare the current sample: in high-evidence frames, 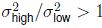, while in low-evidence frames, 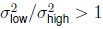. Adopting a more variable belief distribution for the target may seem counterintuitive, but the core idea is that a broader (higher-variance) representation of the goal-relevant category assigns non-negligible likelihood to a wider range of evidence values, meaning ambiguous or extreme samples are more readily attributed to that category. This has an impact on the confidence generation process, producing the kind of overweighting of goal-congruent evidence that characterises the positive evidence bias. This is achieved in the absence of heuristic biases affecting confidence reports — instead, a PEB emerges directly from how belief precision is allocated according to task goals

GOAL model simulations were generated using asymmetric belief variances in the high (*σ*_high_ = 1.2, *σ*_low_ = 1) and low evidence (*σ*_high_ = 1, *σ*_low_ = 1.2) frames, producing goal-dependent choice behaviour and asymmetries in how evidence was integrated into confidence. A logistic regression predicting simulated choices showed that, in the high frame, the option with higher evidence was selected, whereas in the low frame, the option with lower evidence was chosen (Figure 4A). In both frames, simulated confidence displayed a U-shaped pattern relative to the evidence difference between the two options (Figure 4A).

**Figure 4:**
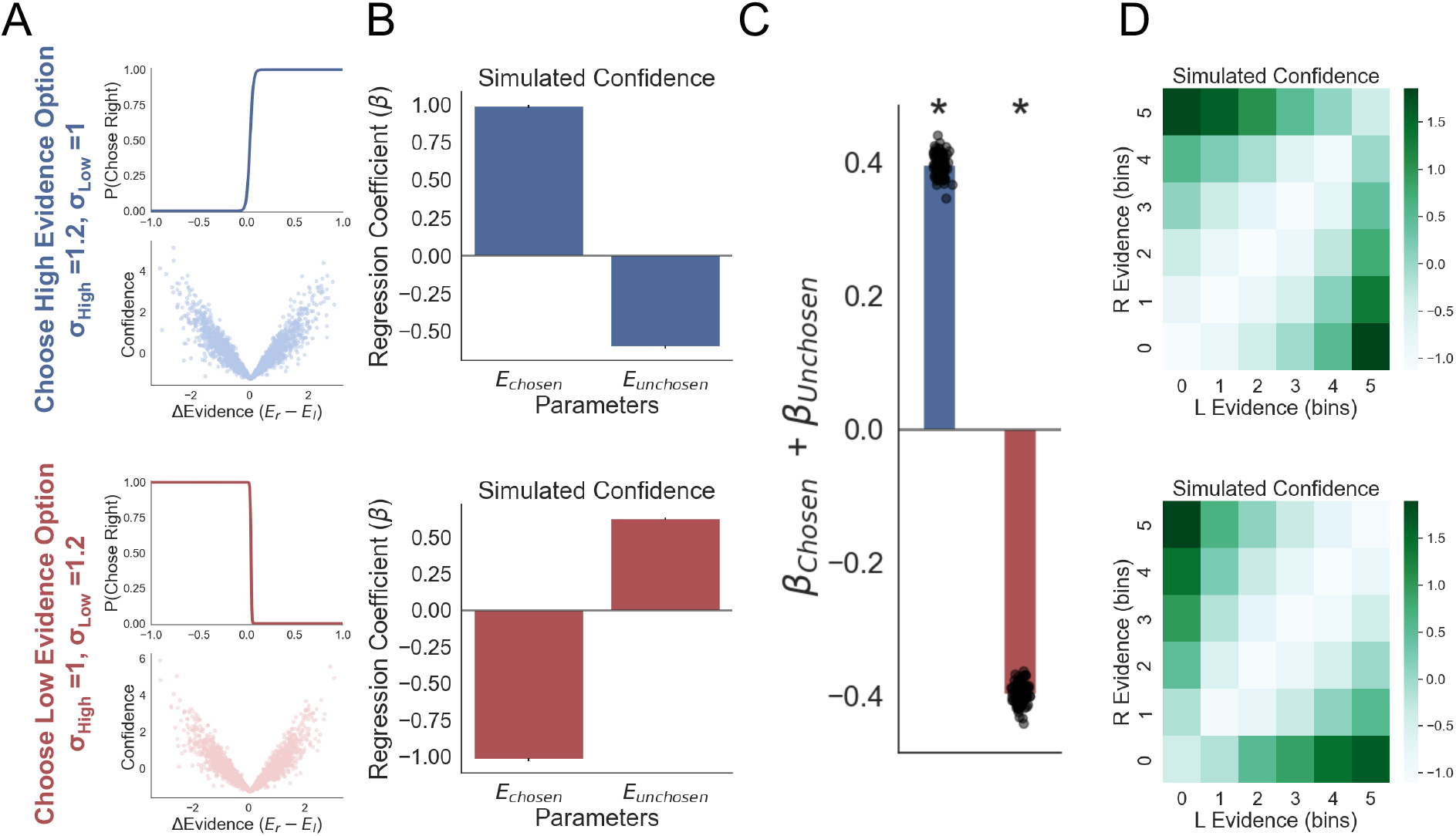
Goal-oriented Asymmetric Likelihood (GOAL) model simulations. High evidence (blue) and low evidence (red) decision frames. (A) Simulated choice and confidence varied with evidence, following standard behavioural patterns. (B) Linear regressions predicting confidence using chosen and unchosen evidence as predictors indicated an overweighting of chosen evidence in both frames. (C) The overall evidence effect (*β*_chosen_ + *β*_unchosen_) from simulated trials reflected the asymmetry in chosen and unchosen evidence integration, with a positive effect of evidence on confidence in the high evidence frame, and a negative effect in the low evidence frame. (D) Confidence in the evidence space, defined by (binned) evidence for left and right options. Displacement of higher confidence levels towards the high evidence values was observed in the high evidence frame. High confidence towards lower values was found in the low-evidence frame.

A linear regression predicting confidence revealed that simulated confidence was mainly driven by the chosen evidence in both frames, with a smaller influence from unchosen evidence (high frame: *β*_chosen_ = 0.994 ± 0.003, *p <* 0.001; *β*_unchosen_ = −0.597 ± 0.003, *p <* 0.001; low frame: *β*_chosen_ = −0.986 ± 0.004, *p <* 0.001; *β*_unchosen_ = 0.606 ± 0.004, *p <* 0.001). As in the human data, the model predicted a positive effect of chosen evidence on confidence in the high frame and a negative effect in the low frame, with the opposite pattern observed for unchosen evidence (Figure 4B).

The GOAL model also reproduced the overall evidence effect (*β*_chosen_ + *β*_unchosen_) and its interaction with task frame (Figure 4C). Overall evidence positively affected confidence in the high frame but negatively in the low frame (high frame: *β*_overall_ = 0.396 ± 0.0016, *p <* 0.001; low frame: *β*_overall_ = −0.396 ± 0.002, *p <* 0.001). The evidence-space plot showed a frame-dependent displacement of confidence for the GOAL simulated confidence reports (Figure 4D). Lower confidence values appeared in the bottom-left quadrant of the high frame and the top-right quadrant of the low frame, consistent with a goal-dependent modulation of confidence.

Finally, we confirmed that similar patterns were observed across different combinations of *σ*_high_ and *σ*_low_, provided that the variance asymmetry followed a goal-dependent rule, with higher variance for the goal-relevant distribution in each task (Supplemental Figure S1).

#### 2.2.3 Heuristic model

Finally, we simulated trials from a model that considers a metacognitive heuristic in which only evidence supporting the chosen option is used to generate confidence.^10,35,42^ In our modelling, the heuristic model (HM) shares a similar architecture with the EVM model at the choice level, which allows replicating the frame-dependent flip in choice behaviour (Figure 5A). However, since in this model confidence is calculated differently, and generated directly from the chosen evidence, HM predicts an inverted U-shape for confidence in the low frame (Figure 5A). This means confidence in the negative frame was lower when one of the alternatives had much lower evidence than the other, which occurs on the extremes of the ΔEvidence axis. In other words, when evidence for the chosen option was weak in the low evidence frame, this model predicts low confidence. Participants did not show this pattern, since they displayed higher confidence when lower evidence was selected in the negative frames, as shown in our previous analysis. Notice that in the EVM and GOAL models, confidence is extracted from the likelihood ratio, i.e., how likely a decision was correct considering both the evidence and statistical features of the belief distributions. Confidence in HM depends *exclusively* on the chosen evidence (high frame: *β*_chosen_ = 1, *p <* 0.001, *β*_unchosen_ = −1.926 × 10*^−^*^16^ ≈ 0, *p <* 0.001; low frame: *β*_chosen_ = 1, *p <* 0.001; *β*_unchosen_ = 1.743 × 10*^−^*^17^ ≈ 0, *p <* 0.001). Unlike human data, the effect of chosen evidence on confidence was always positive, disregarding the change in the frame (Figure 5B). In line with these results, we found the simulations with HM could replicate the overall effect of evidence on confidence, but not the flip of the effect in the low frame (high frame: *β*_overall_ = 1, *p <* 0.001; low frame: *β*_overall_ = 1, *p <* 0.001) (Figure 5C). The bidimensional evidence plot shows how the heuristic definition does not account for the changes in confidence when the goal of the task shifts (Figure 5D). It could be argued we are presenting an unfair representation of the heuristic model, which could be easily adjusted to capture the confidence effects in the low evidence frames, for example, by adapting evidence at the input so the low evidence cases become relevant in the negative frames.^15^ However, implementing this model variant means that the core rationale motivating the heuristic model would no longer hold (the direct mapping between stimulus strength and confidence), and further modifications would become increasingly ad hoc.

**Figure 5:**
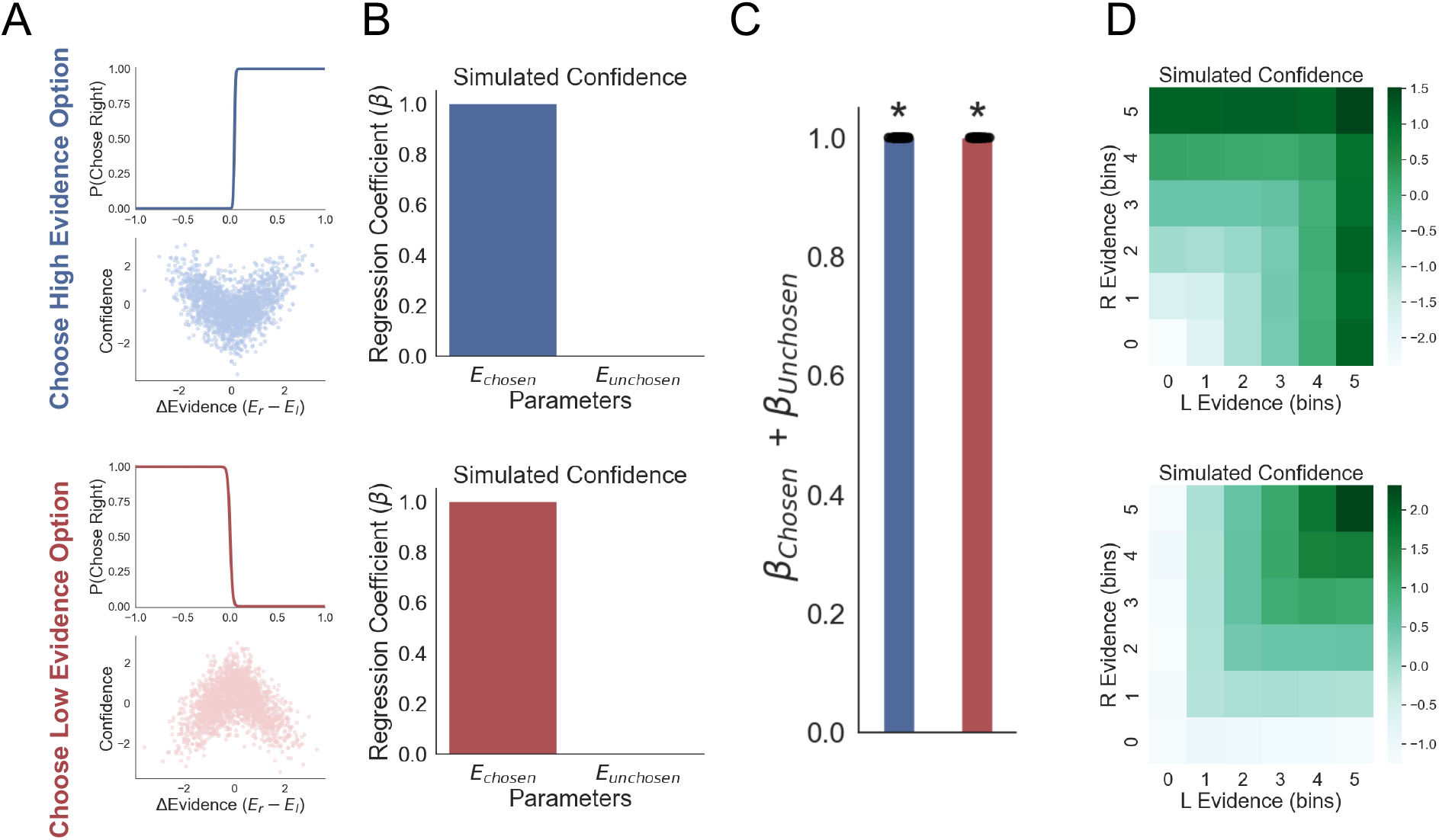
Heuristic Model (HM) simulations. Frames in which the model selects high evidence (blue) or low evidence (red) are presented. (A) Simulated choice and confidence. HM fails to simulate a standard U-shape for confidence in the low evidence frame. (B) Linear regressions predicting confidence using chosen and unchosen evidence as predictors indicated that only the chosen option contributes to confidence. (C) The overall evidence effect (*β*_chosen_ + *β*_unchosen_) from simulated trials reflected the integration of chosen over unchosen evidence, yet the frame did not affect the sign of this integration. (D) Confidence in the evidence space, defined by (binned) evidence for left and right options.

### 2.3 Fitting the GOAL model to human data

In the previous section, we showed that the GOAL model generates a goal-dependent evidence bias on confidence as observed in human behaviour. In this model, the asymmetry of belief variance is crucial to generate this effect. In the following analysis, we asked whether the model fitted to experimental data is sufficient to generate variance asymmetries that follow the same patterns predicted by the simulations. The generation of confidence reports in this version of the model also incorporated reaction time (RT) information, such that the fitting process controls for its influence. Model fitting was performed in a Bayesian framework, pooling together all the even-numbered trials across participants. The fitted model was evaluated on the odd-numbered trials. Samples from the posterior distribution of fitted parameters were used to generate synthetic choices and confidence (Simulations). Analysis of participant’s behaviour (Human) was presented as a reference (Figure 6). We replicated the patterns of results observed in human participants; however, we are not making any claim that the magnitude of the effects in simulations is comparable with the magnitude of human regression parameters.

**Figure 6:**
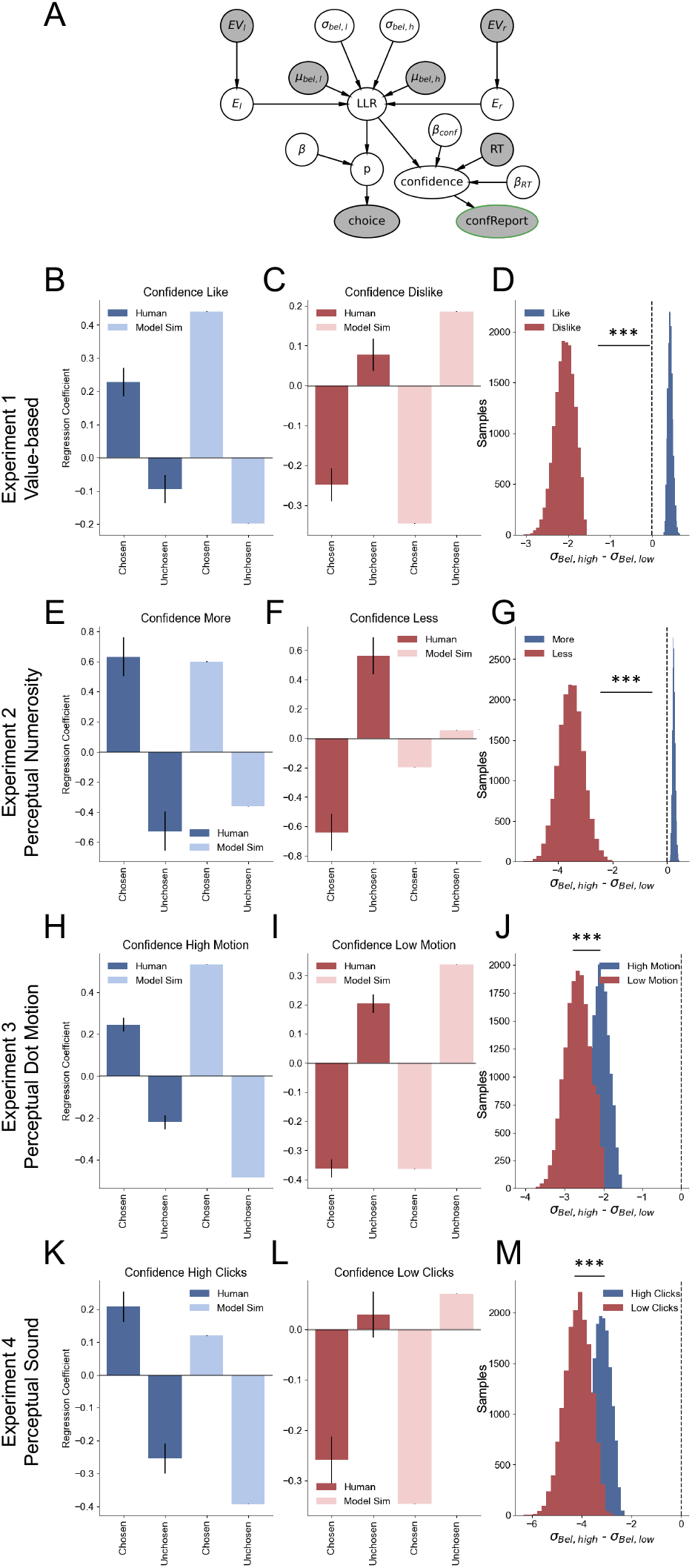
GOAL model fit to decision making experiments. (A) Bayesian graphical model indicating the integration of evidence to generate choice and confidence reports. Shaded circles indicate observed variables. Regression analysis predicting confidence for human data and model simulations. Asymmetries in the value of the latent parameters, belief variance (*σ*_bel_), were found in the model fitted to participants’ data. *EV_l_* and *EV_r_* are the evidence values used in the experiments. *E_l_* and *E_r_* are trial-wise evidence estimates for the left and right options. *µ_bel,l_*and *µ_bel,h_* are the fixed means of the low– and high-evidence generative distributions. *LLR* is the log-likelihood ratio comparing the evidence for the right vs. left option under the low– and high-evidence distributions. *β* is a free parameter scaling the influence of *LLR* on choice probability. *p* is the trial-wise probability of choosing the right-side option. *β_conf_* and *β_RT_* are free parameters scaling the influence of *LLR* and reaction time, respectively, on predicted confidence; *RT* is the observed reaction time for each trial. *LLR*, *RT*, *β_conf_*, and *β_RT_* are used to generate the overall confidence variable (*confidence*), which is transformed into the noisy confidence report (*confReport*). (B–D) Experiment 1: model simulations for like (B) and dislike (C) frames. Simulations replicate the pattern of results obtained from the regression analysis on participants’ behaviour. Specifically, the model captures the frame-dependent variation of the chosen and unchosen evidence effect on confidence. Latent parameters, *σ*_high_ and *σ*_low_ are distinct depending on the frame, with the variance of the goal-relevant alternative relatively higher (D). This means that *σ*_high_ is higher in the like frame and *σ*_low_ is higher in the dislike frame. (E–G) Experiment 2: model simulations for *most* (E) and *fewest* (F) frames, replicate the behavioural effect of chosen and unchosen on confidence, and its interaction with the frame. Belief variance also presents the expected asymmetry, with higher variance for the goal-relevant option (G). (H–J) Experiment 3: model simulations for *high* (H) and *low* dot motion frames (I). Simulations replicate human behaviour in confidence. The values of *σ*_bel_ for high and low distributions were significantly different between frames (J). (K–M) Experiment 4: The model simulations captured the pattern of confidence effects for the auditory experiment in the *high* (K) and *low* (L) clicks frames. The variance distributions *σ*_bel_ for *high* and *low* clicks were also significantly different depending on the frame. Linear regression models are presented for confidence in participants and model simulations (pooled linear regression). *σ*_bel,high_ – *σ*_bel,low_ distributions in panels were generated from sampling posterior distributions of the fitted models.

#### 2.3.1 Experiment 1

The GOAL model fitted to the value-based experiment’s data replicated goal-dependent changes in behaviour. We found that in simulated choices in the *like* frame, the high value item was preferentially selected, while in the *dislike* frame the low value item was picked, an effect captured by a logistic model (logistic function slope, *β*_like_ = 0.872, *β*_dislike_ = −0.751). From a linear regression model predicting confidence, we found the simulated trials also displayed a goal-dependent bias effect (Figure 6B–C), with an increase in confidence during the *like* frame when the chosen item had higher value, and a decrease of confidence in the *dislike* frame (*like*: *β*_chosen_ = 0.440 ± 0.001, *p <* 0.001; *dislike*: *β*_chosen_ = −0.347 ± 0.001, *p <* 0.001). In both frames, the effect of chosen evidence on confidence was greater than that of unchosen evidence (*like*: *β*_unchosen_ = −0.199 ± 0.001, *p <* 0.001; *dislike*: *β*_unchosen_ = 0.186 ± 0.001, *p <* 0.001). The model predicted an asymmetry in the latent variables *σ*_high_ and *σ*_low_ (Figure 6D), with the difference of the belief variances (Δ*σ*_bel_ = *σ*_high_ − *σ*_low_) indicating that the high value distribution had higher variance in the *like* frame (mean *σ*_low_ = 5.177; mean *σ*_high_ = 5.611; mean Δ*σ*_bel_ = 0.435, *p <* 0.001) and the low value distribution presented higher variance in the *dislike* frame (mean *σ*_low_ = 6.345; mean *σ*_high_ = 4.267; mean Δ*σ*_bel_ = −2.078, *p <* 0.001). The value Δ*σ*_bel_ was significantly different between *like* and *dislike* frames (*like* Δ*σ*_bel_ – *dislike* Δ*σ*_bel_; *t* = −1305.54, *p <* 0.001), indicating the asymmetries in variance were distinct for each decision goal.

#### 2.3.2 Experiment 2

In the perceptual experiment on dot numerosity, the GOAL model was also successful in capturing participants’ behaviour. The model predicted frame-dependent choice behaviour (logistic function slope, *β*_most_ = 3.791, *β*_few_ = −1.049). The simulations for this experiment also captured a goal-dependent evidence bias on confidence, with a boost of confidence when the number of dots in the chosen option was higher in the *most* frame (most: *β*_chosen_ = 0.599 ± 0.001, *p <* 0.001) (Figure 6E) and an increase in confidence when the number of dots in the chosen option was lower in *fewest* frame (*fewest*: *β*_chosen_ = −0.197 ± 0.001, *p <* 0.001) (Figure 6F). The model also captured that chosen evidence had a greater effect over confidence than unchosen evidence (*most*: *β*_unchosen_ = −0.365 ± 0.001, *p <* 0.001; *fewest*: *β*_unchosen_ = 0.056 ± 0.001, *p <* 0.001). In this experiment the latent variables of the GOAL model, *σ*_high_ and *σ*_low_, also presented asymmetries depending on the frame (Figure 6G): in the *most* frame the distribution characterising higher evidence (number of dots) presented higher variance (mean *σ*_low_ = 4.763; mean *σ*_high_ = 5.003; mean Δ*σ*_bel_ = 0.240, *p <* 0.001), while in the *fewest* frame the low evidence distribution was found to have the higher variance (mean *σ*_low_ = 6.502; mean *σ*_high_ = 2.972; mean Δ*σ*_bel_ = −3.530, *p <* 0.001). Similarly to Experiment 1, the two frames presented a distinct balance between *σ*_high_ and *σ*_low_ (*most* Δ*σ*_bel_ – *fewest* 401 Δ*σ*_bel_; *t* = −644.25, *p* < 0.001).

#### 2.3.3 Experiment 3

We again found a similar pattern of results for the simulated trials using the GOAL model fitted to our dot motion experiment. From the logistic regression analysis, we found that synthetic choices selected high motion and low motion directions depending on the trial frame (logistic function slope, *β*_highMotion_ = 2.250, *β*_lowMotion_ = −1.988). The simulated effects of chosen and unchosen evidence on confidence resembled the values estimated from human responses. Chosen evidence had a positive and negative influence on confidence in the *high and low motion* frames, respectively (*high motion*: *β*_chosen_ = 0.5332 ± 0.001, *p <* 0.001; *low motion*: *β*_chosen_ = −0.364 ± 0.001, *p <* 0.001). Similarly to previous experiments, the influence of unchosen evidence on confidence was comparatively smaller than that of chosen evidence (*high motion*: *β*_unchosen_ = −0.483 ± 0.001, *p <* 0.001; *low motion*: *β*_unchosen_ = 0.339 ± 0.001, *p <* 0.001) (Figure 6H–I). These results indicate that the model captured the goal-dependent evidence bias. However, unlike Experiments 1 and 2, the asymmetry of belief variance (Figure 6J) was negative in both frames: the low evidence distribution had a higher variance in the *high motion* frame (mean *σ*_low_ = 6.057; mean *σ*_high_ = 3.964; mean Δ*σ*_bel_ = −2.093, *p <* 0.001) and the *low motion* frame (mean *σ*_low_ = 6.259; mean *σ*_high_ = 3.588; mean Δ*σ*_bel_ = −2.671, *p <* 0.001). We attribute this to the parametrisation of our model, since to employ an identical model in relation to our previous experiments, we used as evidence inputs the number of dots, when in other experiments coherence or directionality are used as evidence inputs (e.g., fleming2018). Despite these discrepancies, the two frames presented a distinct balance between *σ*_high_ and *σ*_low_ (*high motion* Δ*σ*_bel_ – *low motion* Δ*σ*_bel_; *t* = −189.79, *p <* 0.001), with a relatively higher variance of the low evidence distribution in the *low motion* frame, in line with the goal of the task.

#### 2.3.4 Experiment 4

Finally, we found that the GOAL model could capture the behavioural signatures we observed in the sound discrimination experiment. From the logistic regression analysis, we found that simulated choices selected high clicks and low clicks directions depending on the trial frame (logistic function slope, *β*_highclicks_ = 1.306, *β*_lowclicks_ = −1.160). The simulations of confidence captured the effect of chosen evidence observed in participants, with a positive effect of chosen evidence on confidence in the *high clicks* frame, while chosen evidence had a negative effect on the *low clicks* frame, similarly to previous experiments (*high clicks*: *β*_chosen_ = 0.120 ± 0.001, *p <* 0.001; low clicks: *β*_chosen_ = −0.346 ± 0.001, *p <* 0.001) (Figure 6K–L). The unchosen evidence had an opposite effect on confidence in both frames (*high clicks*: *β*_unchosen_ = −0.394 ± 0.001, *p <* 0.001; *low clicks*: *β*_unchosen_ = 0.07 ± 0.001, *p <* 0.001). In this case, the model predicts a stronger influence of unchosen evidence over confidence, in comparison to chosen evidence, which mirrors the behaviour that we observed in human responses. The asymmetry of belief variance was also observed in this experiment (Figure 6M), although the model predicted a higher value of the low evidence distribution in both frames (*high clicks*: mean *σ*_low_ = 6.598, mean *σ*_high_ = 3.366; mean Δ*σ*_bel_ = −3.231, *p <* 0.001; *low clicks*: mean *σ*_low_ = 6.772; mean *σ*_high_ = 2.597; mean Δ*σ*_bel_ = −4.176, *p <* 0.001), in line with the stronger influence of negative effects on confidence (unchosen evidence in *high clicks* and chosen evidence in *low clicks*). Still, the two frames presented a different balance between *σ*_high_ and *σ*_low_ (*high clicks* Δ*σ*_bel_ – *low clicks* Δ*σ*_bel_; *t* = −153.01, *p <* 0.001), indicating an impact of task goal on the generation of confidence.

Overall, these results show that our GOAL model can replicate goal-dependent evidence biases on confidence across a spectrum of decision tasks, hinting that a modulation of the variance of latent variables could provide a unifying explanation of this phenomenon in human perception and decision making.

### 2.4 Attention as a cognitive mechanism for the PEB

Our GOAL model is agnostic to the cognitive process that drives the asymmetry in variance and triggers the effect on confidence. One hypothesis is that selective attention might underpin this process, generating an imbalance in information processing. In previous work,^15^ we reported that visual attention is preferentially allocated towards the alternatives that have higher relevance for the goal of the task. This was found in the original analysis of the data reported here in Experiments 1 and 2, e.g., in value-based and perceptual tasks. Furthermore, in that same work, we reported that gaze-weighted accumulator models^43^ could capture the overall evidence effect when estimating confidence from a balance of evidence model.^44^ However, this behaviour was captured only if information about attentional allocation was used by the model. These results, in line with our current SDT-inspired model, hint that goal-directed asymmetries in the integration of evidence for a decision are key to explaining confidence.

In this section, we further investigated the hypothesis that goal-dependent effects of attention can explain biases in confidence. The results of our Experiment 3 give a hint of this effect: in the dot motion experiment with a single aperture, the goal-relevant evidence bias in confidence was present even when the spatial displacement of attention was not required. This finding implies a more general attentional process. In the fourth experiment, we further expanded this to the auditory dimension to isolate the visual attention process in a task using dichotic click sequences, while pupil size variation was tracked.^25^ In the *High Clicks* frame, participants were instructed to select the side with more clicks; in the *Low Clicks* frame, they were instructed to select the side with the lower number of clicks (Figure 1G).

As reported above for Experiment 4 (Figure 1H), we observed goal-dependent asymmetries in the effect of evidence on confidence reports. We further analysed the variation in relative pupil traces in each trial (please see Methods for details). We found that after the presentation of the sound stimuli, pupil size increased, reaching a peak around choice time, following a pattern similar to that observed in other decision experiments.^25,28^ Overall, trials reported with low confidence had on average higher pupil size at decision time (average relative pupil area, high confidence = 0.79 ± 0.07; low confidence = 0.88 ± 0.08, *t*(31) = −3.08, *p <* 0.01). Trials with higher difficulty (i.e., lower differences in the number of clicks between the options) also presented higher pupil sizes relative to easier trials at decision time (average relative pupil area, high |ΔClicks| = 0.81 ± 0.07; low |ΔClicks| = 0.86 ± 0.07, *t*(31) = −2.34, *p <* 0.05). Correct trials also had on average lower pupil size during the decision period compared with error trials (average relative pupil area, correct = 0.82 ± 0.07; incorrect = 0.91 ± 0.09, *t*(31) = −3.09, *p <* 0.01).

Since we found that trial difficulty and confidence had an effect on pupil size at decision time, we centred our analysis on this period, separately for each frame. This focus on the pre-decision time window characterises the deliberation stage, where we expect to find asymmetries in evidence integration. We focused our analysis on the effect of chosen evidence on pupil variation since the results above highlighted the relevance of this factor on confidence. We separated trials using a median split of the number of clicks in the chosen option (Figure 7B). We found that when participants’ goal was to choose the higher evidence alternative (*High Clicks* frame), their pupil area was smaller in trials with a higher number of clicks. Conversely, in the *Low Clicks* frame, their pupil size was reduced in trials with a lower number of clicks. In other words, the frame affected how the pupil responded to the chosen evidence. To corroborate this finding, we fitted a linear regression model predicting relative pupil size at each time point at the individual participant level. We included chosen and unchosen evidence as predictors, in addition to reaction time and pupil position in the *X*–*Y* coordinates.^28^ We found that during the pre-decision period, chosen evidence had a negative effect on pupil size in the *High Clicks* frame, while this effect was positive in the *Low Clicks* frame (Figure 7C). These regression coefficients were significantly different between frames around 1 second before the choice (average regression coefficient for the time points with a significant difference between frames: *β*_HighClicks_ = −0.052 ± 0.005, *β*_LowClicks_ = 0.033 ± 0.009, Δ*β* FDR-corrected permutation test *p <* 0.01, cluster size = 6, indicated with black line in Figure 7C). This confirms that in the positive frame, an increase in chosen evidence reduces the pupil size, while in the negative frame, an increase in chosen evidence boosts pupil size.

**Figure 7:**
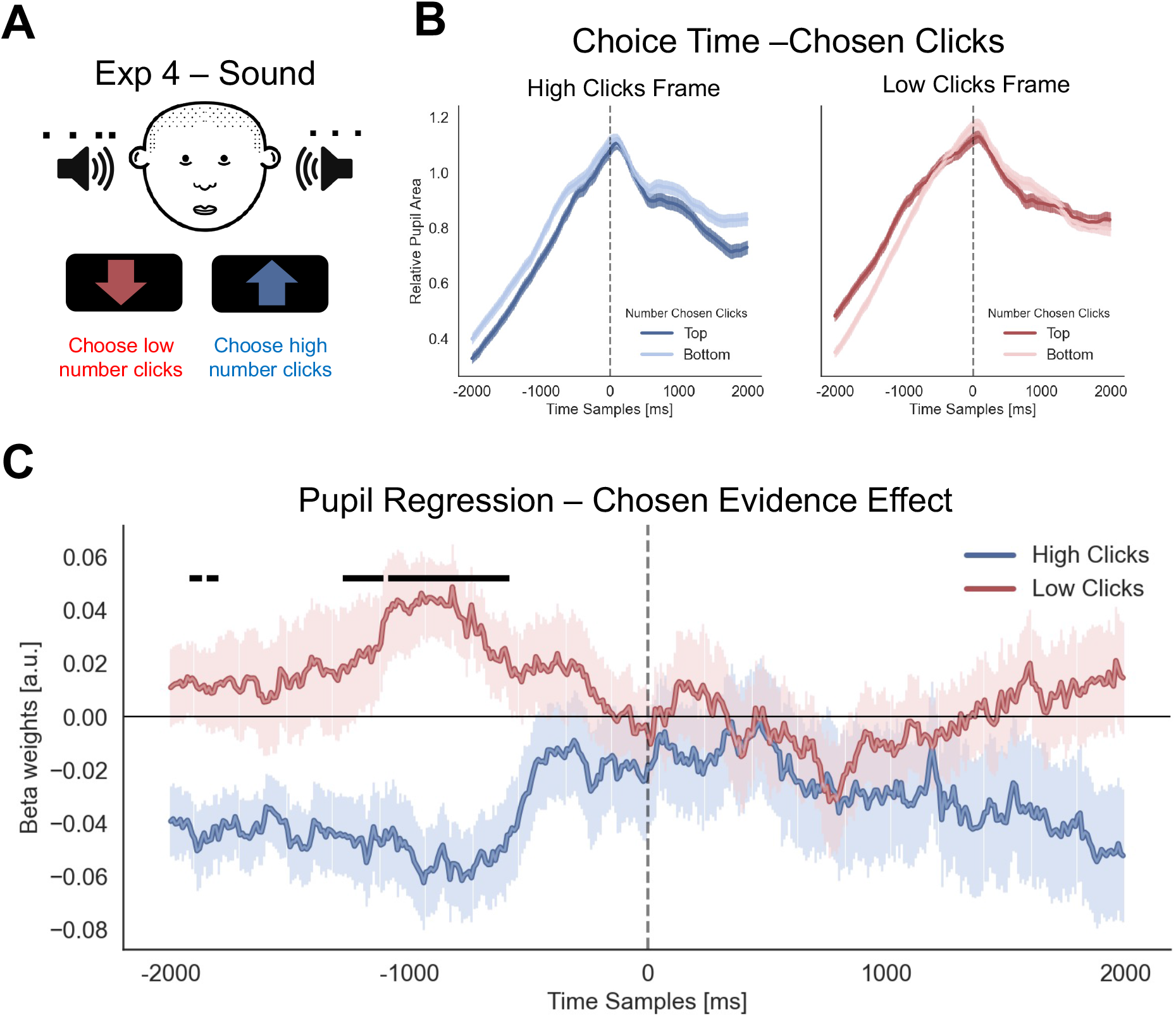
Pupil variations depend on the task goal. (A) Auditory decision experimental design. In each trial, sequences of click sounds were presented to left and right-side speakers and participants were instructed to select the option with a higher or lower number of clicks, depending on the frame. Participants were free to respond at any time. They reported their confidence in the decision at the end of each trial. Pupillometry was tracked throughout the whole process. (B) Pupil traces by frame. Trials were separated using the median split of the number of clicks in the chosen alternative. For the pre-decision time, while in the High Clicks frame, participants showed lower pupil size in the trials with a higher number of clicks; in the Low Clicks frame the participants had a lower pupil size when they chose the most appropriate evidence for that trial, the option with fewer clicks. Lines represent average pupil trace across all participants’ trials. The shaded area indicates standard error. (C) We corroborated the pupil results with a linear regression analysis predicting relative pupil area at each time point. The effect of chosen clicks on the pupil is presented in the figure. During the pre-decision time, a significant difference in the effect of chosen clicks was found between High and Low Clicks frames. Lines represent the average effect of the chosen evidence parameter estimated at a participant level. The shaded area indicates the standard error, estimated from the participants’ *β* group sample. The black line indicates a significant difference between frames, estimated from an FDR-corrected permutation test.

Overall, we show that an increase in pupil size is most prominent on trials in which evidence diverged from the participants’ goal. We suggest that the change in frame triggers attentional reorienting aimed at gathering the relevant information for the task, reflected by both the goal-dependent change of priors in our model and the dynamics of pupil size.

## 3 Discussion

Our results show that confidence is formed relative to the goals of the decision-maker: when the goal changes, the mapping between evidence and confidence reverses, across perceptual and value-based domains alike. The clearest demonstration of this principle is our observation of a “flip” in the positive evidence bias (PEB) which has long been treated as a fixed feature of the confidence computation.^8–10,42^ One influential account holds that the bias arises because, in discrimination tasks, the decision system operates in a detection-like mode, treating the problem as one of identifying whether a stimulus is physically present rather than comparing two competing options.^10,18,19^ Detection naturally favours the option carrying more evidence, a pattern captured by heuristic models that compute confidence solely from the chosen option and discard the unchosen one. Virtually all studies of the PEB, however, employ tasks with a single goal: selecting the option with the stronger evidence. Under these conditions, positive evidence and task goal are perfectly confounded. Our frame manipulation breaks this confound by requiring participants to sometimes select the option with less evidence. If the PEB were purely stimulus-driven, it should have remained positive regardless of frame. Instead, it reversed: in negative frames, weaker evidence for the chosen option increased confidence. Crucially, this reversal emerged not only in perceptual tasks but also in value-based choices, where no physical signal is available to be detected. Together, these findings argue against accounts of confidence formation rooted in low-level stimulus encoding, and point instead to a domain-general mechanism by which goals determine how evidence is weighted in the construction of confidence. In other words, confidence tracks goal-fulfilment, not stimulus evidence or decision accuracy.

Across all four experiments, choice accuracy is preserved across frames, but confidence shifts systematically with the goal-relevant dimension of the stimulus. Standard accounts treat confidence as a probabilistic readout of decision accuracy,^1^ under which confidence and accuracy should rise and fall together, and cannot accommodate this dissociation. What confidence appears to track, instead, is the degree to which the chosen option exemplifies what the agent was looking for: how strongly it instantiates the goal-relevant property. For example, in the *most* frame, more dots in the chosen patch increase confidence; in the *fewest* frame, fewer dots do. Both reflect the same underlying signal, where the chosen option is a clearer instance of the category the goal has made relevant. Framed this way, confidence is less a verdict on whether the decision was correct and more a readout of how well the choice satisfies the current goal. This interpretation is aligned with the idea of confidence as a measure of self-consistency^45–48^ as observed in value-based or aesthetic choices.^36,49^ This functional role is well suited to real-world decision-making, where confidence supports persistence, calibrates learning, and is communicated to others—all of which require a signal calibrated to goal-fulfilment rather than to abstract accuracy. This view aligns with proposals that metacognition functions as a control system rather than a purely epistemic readout.^50,51^

To formalise this account, we developed GOAL (Goal-Oriented Asymmetric Likelihood), a Bayesian observer model grounded in signal detection theory. Like other ideal observer models,^38^ GOAL assumes the agent has internalised the task structure through two belief distributions, representing high-evidence and low-evidence options respectively. While standard SDT models assume equal variance,^10,34,52^ GOAL allows the variances to differ, with the goal-relevant distribution assigned higher variance. The observer computes the likelihood ratio that the two options were drawn from these distributions, and uses its magnitude both to choose and to generate confidence. This single modification reproduces the full pattern of human confidence reports: chosen-evidence overweighting, ΣEvidence modulation, and the reversal of both effects when the goal shifts. Mechanistically, the variance asymmetry implements a goal-conditioned selection of which evidence dimension dominates the confidence computation, ensuring the chosen option’s evidence is always overweighted, while the readout dimension is set by the goal.

GOAL contrasts with two existing classes of model. Equal-variance balance-of-evidence models, which represent decision accuracy alone, cannot capture chosen-evidence overweighting in either direction.^8,10,15,35^ Heuristic models that compute confidence from chosen evidence alone^10,18,19^ capture the bias and the ΣEvidence effect, but cannot easily accommodate frame reversals. Our data also show that unchosen evidence is not fully discarded: it contributes systematically, with opposite sign, to confidence. Recent extensions allow flexible weighting of unchosen evidence;^53^ whether such models accommodate goal-dependent reversals remains to be tested.

A recent proposal suggests that detection-like confidence and PEB can emerge from computing confidence in a high-dimensional evidence space, where the goal entertained by the subject determines which dimension is selected as the target.^20^ This Bayesian framework predicts that variance asymmetries in confidence can also reflect a goal-dependent “flip” in evidence bias, by reassigning the target when subjects are asked to report the lower value item in a pair. However, Kozyra et al. still assume equal-variance evidence representations by construction, locating the asymmetry entirely in the confidence computation itself. GOAL instead places the asymmetry in the evidence representation. These accounts are not mutually exclusive, as dimensionality-driven normalisation and representational variance asymmetries could jointly contribute to the observed reversals, and their relative contributions remain to be tested empirically.

A natural candidate mechanism for the goal-conditioned weighting in GOAL is selective attention. We previously showed that visual attention is preferentially allocated to goal-relevant alternatives,^15^ and related work has proposed that attention modulates the reliability of accumulated evidence.^54,55^ Crucially, our dot-motion and auditory experiments show that the effect does not require spatial displacement of attention, as the same reversal emerges when all evidence appears at a single location or in a non-spatial modality. We therefore propose a more general attentional gain on goal-relevant information, expressed as increased receptivity to evidence in the goal-relevant category. This places the evidence bias within the decision process itself,^20,56^ consistent with reports that the PEB does not depend on noisy encoding^17^ or post-decisional processing.^13^

Our pupillometry results provide independent, physiological support for goal-conditioned asymmetries in evidence integration: pupil size was lower on goal-congruent trials, suggesting that goal-relevant evidence elicited less autonomic arousal at decision time. This pattern is consistent with the goal acting as a prior against which incoming evidence is evaluated, drawing loosely on prediction-error accounts linking pupil dilation to the mismatch between expectations and outcomes.^27,30,57^ The more strongly an expectation is held, the greater the surprise when it is violated. In our task, the frame sets this expectation. In the *High Clicks* frame, participants expect the higher-evidence option to satisfy their goal, so choosing an option with relatively more clicks is less surprising than choosing one with fewer, even when that choice is correct. The *Low Clicks* frame reverses this expectation, producing the corresponding flip in pupil response.

Some limitations are worth noting. The auditory experiment showed a negative overall effect of ΣEvidence in the positive frame, opposite to the visual experiments. This is likely a consequence of the fixed stimulus window, though the goal-dependent modulation by frame was preserved. The simultaneous presentation of both auditory streams also prevented us from testing GOAL’s prediction that variance is specifically increased for the goal-relevant alternative; sequential designs would address this directly. Future work should also examine the impact of goal-tuning on metacognitive sensitivity,^38^ which our frame manipulation is well-suited to investigate.

In conclusion, our results suggest that confidence does not simply track decision accuracy or stimulus evidence, but instead monitors how well a choice fulfils the decision-maker’s goals. This conclusion extends well beyond the PEB, which emerges here as one signature of a general goal-relative computation, one that previous tasks could not reveal because the goal was always tied to the stronger option. Given that this effect reflects an adaptive adjustment to the demands of the current goal, characterising it as a bias may be inaccurate. What appears as a distortion in goal-neutral laboratory tasks may instead reflect the normal operation of confidence: a signal that tracks how well a choice serves the goals of the decision-maker.

## Methods

### M.1 Procedure

Across four experiments, we tested how goal-dependent framing shapes decision-making and confidence in binary choice tasks in value-based and perceptual domains. In each experiment, participants chose between two simultaneously presented options and then rated their confidence in that choice. Critically, every experiment employed a within-subject framing manipulation, alternating in blocks, in which participants were instructed to select either the option with more evidence of an attribute (e.g., preferred value, dot numerosity, coherent motion, or auditory clicks) or the option with lower evidence of that attribute. The four experiments varied systematically in stimulus domain and modality: Experiment 1 used snack-food subjective valuations in a value-based choice task; Experiment 2 used visual dot-numerosity arrays; Experiment 3 used random dot motion kinematograms; and Experiment 4 used dichotic auditory click trains.

#### M.1.1 Experiment 1 – Value-based decision

Experiments 1 and 2 use datasets previously reported in.^15^

At the start of the session, participants indicated how much they were willing to pay (from £0 to £3) for each of 60 snack food items. This valuation phase followed the Becker–DeGroot–Marschak (BDM) procedure,^32^ which incentivises truthful reporting by linking participants’ stated bids to the possibility of actually purchasing one of the items at the end of the experiment. Participants were asked to fast for four hours before the session, in line with established protocols for value-based decision studies.^14,58,59^

Following the bidding phase, participants completed a binary choice task. On each trial, two snack items appeared on the screen—one to the left and one to the right of centre (Figure 1). Participants selected one of the two items and then rated their confidence in that choice. Pairs were created based on individual value ratings: using a median split, each item was classified as high-value or low-value, generating 15 high-value, 15 low-value, and 30 mixed pairs (60 pairs total), tailored to each participant’s preferences. Each pair appeared twice, with left–right positions reversed to counterbalance presentation order.

A key feature of the design was the use of two framing conditions: (1) *Like frame*: participants chose the item they preferred to eat. (2) *Dislike frame*: participants chose the item they liked least, knowing this meant they would receive the other item for consumption at the end.

After four practice trials, participants completed six blocks of 40 trials each (240 trials total). Like and dislike frames alternated between blocks, and the starting frame was counterbalanced across participants (120 trials per frame). A small icon in the top-left corner of the screen (“thumbs up” for like, “stop sign” for dislike) reminded participants of the current task frame, and the experimenter announced the frame at the start of each block. The final trial of one block was never the first of the next.

Eye movements were recorded throughout the task. Stimuli were presented using a gaze-contingent design so that participants viewed only one item at a time, depending on where they looked. This prevented them from processing both options simultaneously in peripheral vision.

At the end of the experiment, one trial was randomly selected for the BDM procedure.^32^ The bid associated with the relevant item (the chosen item in the like frame, or the unchosen item in the dislike frame) was compared with a randomly generated price between £0 and £3. If the participant’s bid exceeded this price, the corresponding amount was deducted from their £20 compensation, and they received the snack item. If the bid was lower, they kept the full £20 and did not receive a snack. In both cases, participants remained in the testing room for one hour afterwards and could not eat anything other than the item purchased through the auction. The entire procedure was explained in advance to ensure full understanding.

#### M.1.2 Experiment 2 – Perceptual decision (dot numerosity task)

Experiment 2 followed a design similar to Experiment 1 but replaced food items with visual stimuli. Participants chose between two circles filled with dots, completing two framing conditions: in the *most* frame, they selected the circle with more dots, and in the *fewest* frame, they selected the circle with fewer dots.

Each trial displayed two dot arrays with total numerosities of 50, 80, or 110 dots. Within each numerosity level, the dot difference between the two circles varied across ten difficulty levels (2% to 20%, in 2% increments). To increase task difficulty, distractor dots (orange) were added alongside the target dots (blue–green), amounting to 80% of the target count (i.e., 40, 64, and 88 distractors for the three numerosity levels). Each pair of dot patches was shown twice, with left–right positions counterbalanced.

After 40 practice trials (20 with feedback and 20 without), participants completed three blocks of 40 trials in the most frame and three in the fewest frame, for a total of 240 trials. Framing conditions alternated between blocks, and the starting frame was counterbalanced across participants. A label (“Most” or “Fewest”) was displayed in the top left corner of the screen to indicate the current frame. After each choice, participants rated their confidence in having made the correct decision.

Stimulus presentation was gaze-contingent, ensuring that only one dot patch was visible at a time, depending on fixation. Participants received £7.50 for the one-hour session. Experiments 1 and 2 were programmed using Experiment Builder (version 2.1.140, SR Research). Although eye-tracking data were collected, they were not used in the present analyses.

#### M.1.3 Experiment 3 – Perceptual decision (random dot motion)

To extend the previous findings, Experiment 3 employed a web-based random dot kinematogram (RDK) task. Participants viewed moving dot displays in which a proportion of dots moved coherently either left or right, while the remaining dots moved randomly. Motion was constrained to horizontal directions to maintain tight control over evidence for and against each choice.

Participants completed two framing conditions: in the *high motion* frame, they selected the direction in which most dots moved; in the *low motion* frame, they selected the direction with fewer dots moving. Displays contained between 20 and 140 dots (seven levels), and each numerosity was combined with eight coherence levels (51%, 52%, 55%, 60%, 65%, 70%, 80%, 90%). Each combination of numerosity, coherence, and motion direction (left/right) yielded unique trials, producing 112 trials per block. Participants completed two blocks per frame (224 trials per frame, 448 in total), with frame order counterbalanced across participants.

Frame type was cued by background colour: blue (#b5cce9) for high motion and red (#e9b5b5) for low motion, matched for brightness. Dots were black, radius 2 pixels, lifetime 20 frames, displayed within an elliptical aperture (500 × 350 px) centred on screen. Trials continued until a response or a 5-second timeout. Participants responded using the A (left) and L (right) keys and rated confidence on a 1–9 scale using number keys.

Before the task, participants completed 32 training trials (with the option to repeat). Performance was checked halfway through the experiment, and participants with accuracy below 60% were excluded. Participants were reimbursed £6 for participation and an additional £4 upon full completion, paid via Prolific. The task was implemented online using the Gorilla platform with jsPsych,^60^ using a modified RDK plugin^61^ that restricted motion to left and right directions.

#### M.1.4 Experiment 4 – Perceptual decision (auditory clicks)

Experiment 4 extended our investigations to the auditory domain using a two-alternative forced-choice task with click stimuli (adapted from^25^). Participants wore headphones and viewed a monitor 1 m away. Each trial presented two 1-second streams of auditory clicks, one to each ear, consisting of brief (40 ms) click bursts on a white noise background. The number of clicks per ear varied from 1 to 10, with all possible unequal combinations used.

Two framing conditions were employed: in the *high click* frame, participants chose the ear with more clicks; in the *low click* frame, they chose the ear with fewer clicks. A central arrow (up or down) indicated the current frame. After each choice, participants rated their confidence on a visual scale.

The experiment comprised four blocks (two per frame) of 45 trials each, totalling 180 trials, with short breaks between blocks. Practice included 20 feedback trials (10 per frame). Each click pair appeared twice, with ear positions counterbalanced. Frame order was randomised across participants.

Participants rested their head on a chin-and-forehead support to stabilise pupil recordings. Each trial began with central fixation (used for drift correction), followed by a 2 s baseline period, then the 1 s auditory stimulus. Participants responded without time limits using the Z (left) and M (right) keys, followed by a 3 s delay and confidence rating using the same keys and spacebar to confirm.

Pupil diameter was recorded at 500 Hz using an EyeLink 1000 eye-tracker (SR Research). The task was programmed in Experiment Builder (version 2.3.38, SR Research) with stimuli generated via custom MATLAB scripts. Clicks were randomly distributed within the 1 s sound window (sampling rate 44.1 kHz). Display resolution was 1024 × 768 px.

### M.2 Participants

#### M.2.1 Experiment 1

Forty volunteers provided informed consent to participate in this study. Data from 31 participants (15 females, 16 males; age range 20–54 years; mean age 28.8) met the inclusion criteria and were retained for analysis. One participant was excluded for failing to use the full value scale during the snack-item bidding task, and another for repeatedly entering identical bid values. Four participants were excluded for giving the same confidence rating on most trials, and three were removed for not following task instructions. Of these, one showed excessive blinking and made selections without fixating on either item, while the other two failed to comply with the frame manipulation. None of the participants reported any current treatment for mental health disorders. To ensure familiarity with the snack stimuli, all participants had lived in the UK for at least one year (mean duration: 17 years).

#### M.2.2 Experiment 2

Forty healthy volunteers were recruited for Experiment 2. Thirty-two participants (22 females, 10 males; age range 19–50 years; mean age 26.0) met the inclusion criteria and were included in the behavioural and regression analyses. Three participants were excluded for giving identical confidence ratings across trials, and five were removed for not following task instructions: four performed near chance level (accuracy below 65%) or ignored the frame manipulation, and one experienced technical issues with eye tracking.

Full details of the exclusion criteria for Experiments 1 and 2 are reported in.^15^ All participants provided written informed consent, and both studies were approved by the University College London Division of Psychology and Language Sciences Ethics Committee.

#### M.2.3 Experiment 3

Experiment 3 was conducted online using the Prolific recruitment platform.^62^ Eligibility criteria included being at least 18 years old, having normal or corrected-to-normal vision, and no diagnosed or ongoing mental health condition. Forty participants were initially recruited; three failed to complete the task and were excluded automatically. Complete datasets were obtained from 37 participants (7 females, 29 males, 1 preferred not to say; age range 18–19 years; mean age 18.2).

After data inspection, three additional participants were excluded for poor performance (accuracy below 65%), four for repetitive confidence ratings (*>*60% of trials with identical ratings), and one for limited use of the confidence scale (*<*50% of its range). The final sample included 29 participants (4 females, 24 males, 1 preferred not to say; mean age 18.2).

All participants completed an online consent form prior to the study. The experiment was approved by the University College London Division of Psychology and Language Sciences Ethics Committee, and data were collected in April 2021.

#### M.2.4 Experiment 4

Experiment 4 was conducted in person using an eye-tracking setup. Thirty-eight participants took part, with 32 retained for analysis (24 females, 8 males; age range 18–52 years; mean age 28.8). Six participants were excluded for poor performance (accuracy below 65%). All participants provided written informed consent before taking part. The study received ethical approval from the University College London Division of Psychology and Language Sciences Ethics Committee, and data were collected between April and October 2022.

### M.3 Data analysis: behavioural data

Behavioural measures during like/dislike, most/fewest, high/low motion, and high/low clicks frames were compared using statistical tests available in SciPy. The Sklearn toolbox in Python was used to perform logistic regressions on choice data. All hierarchical analyses were performed using the lme4 package^63^ for R, integrated into a Jupyter notebook using the rpy2 package (https://rpy2.readthedocs.io/en/latest/). Additionally, we predicted confidence using a linear mixed-effects model. The confidence model included trial-level reaction time (RT), *confidence* ∼ *RT* +*ChosenEv* +*UnchosenEv*. Fixed-effects confidence intervals were calculated by multiplying standard errors by 1.96. Predictors were all z-scored at the participant level. Matplotlib/Seaborn packages were used for visualisation.

### M.4 Bayesian Model

We developed a model inspired by standard two-dimensional signal detection theory (SDT) to describe choice and confidence behaviour in our experiments. In SDT, the alternatives are usually characterised as target and non-target, with two distributions characterising each one of them.^38^ Both target and non-target are modelled using Gaussian distributions with mean evidence (*µ*) and variance (*σ*^2^). In the standard SDT approach, the mean of the target option is larger than the non-target (*µ*_Target_ *> µ*_Nontarget_) and the variance of both distributions is equal (*σ*_Target_ = *σ*_Nontarget_). This framework characterises tasks without modification of goals, i.e., in which the target option always contains more evidence (e.g., number of dots, visual contrast, etc.). In our experiment, target and non-target options were not fixed categories from the perspective of differences in evidence, since different task manipulations demanded choosing options with either higher or lower evidence. To generalise SDT to this case, we designate the target and non-target distributions as high or low evidence, respectively, with *µ*_High_ *> µ*_Low_.

In this section, we describe the details of Bayesian graphical models used for the simulations and model fit to each experiment. All model simulations and fits were implemented in a Bayesian framework using the Python library PyMC.^64^

### M.5 Model Simulations

We simulated a Bayesian observer in a generic binary decision experiment (Figure 2). The agent is exposed to two options, with different amounts of evidence (right and left evidence, *E_r_* and *E_l_*respectively). The simulated agent had to report the option (left or right) with higher evidence in a High Evidence frame and the alternative with lower evidence in a Low Evidence frame. For each simulated trial, we generated the side of the item with higher evidence (side; 0: left; 1: right) from a Bernoulli distribution (High Ev Opt variable in Figure 2B). The evidence for each option was extracted from a Gaussian distribution for the right and left possible directions:

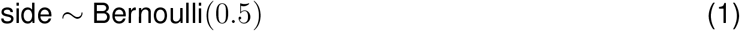

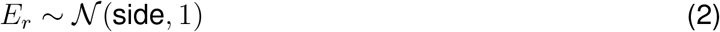

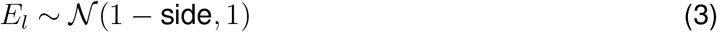

The simulated agent had two internal Gaussian distributions that characterised their beliefs about the distributions of high evidence samples and low evidence samples (Figure 2A), *E*_high_ and *E*_low_, respectively:

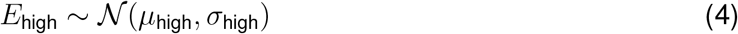

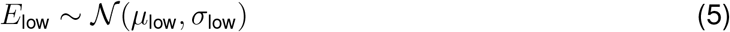

Therefore, for each trial, the simulated agent observed a pair of evidence samples ⟨*E_l_, E_r_*⟩ and determined the likelihood they were generated from the distribution ⟨High Evidence, Low Evidence⟩ or ⟨Low Evidence, High Evidence⟩. The logarithm of the ratio of these two likelihoods (*L*) was used as the decision variable for the observer:

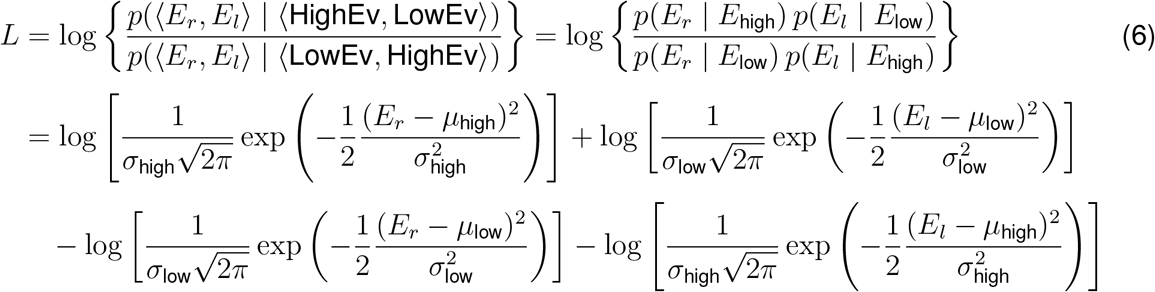

Depending on the frame that was simulated, the observer selected the option with lower or higher evidence. When high evidence was to be picked, the agent chose the right option if *L* ≥ 0 and the left option if *L <* 0. In the low evidence frame, the right option was chosen if *L* ≤ 0 and the left option when *L >* 0. A basic metric of confidence was calculated as the magnitude of the log-likelihood ratio:

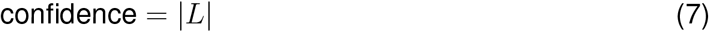

We simulated three different models to study how the generation of confidence is affected by decision parameters. In all these models we assumed that the average values for the high and low evidence distributions was *µ*_High_ = 1 and *µ*_Low_ = 0, respectively.

We simulated a standard equal variance model (EVM) typically assumed in SDT. The choice decision variable (Equation 6) and confidence (Equation 7) were calculated for all the simulated trials. For the simulations in the Results section, we illustrated the model predictions by setting *σ*_high_ = 1 and *σ*_low_ = 1 within the high evidence frame, and *σ*_high_ = 1 and *σ*_low_ = 1 within the low evidence frame (2000 samples were extracted from the posterior distribution). This model approximates a Balance of Evidence model which cannot reproduce the goal-dependent evidence effects on confidence (i.e., no overweighting of chosen evidence).

During exploration of the data patterns, we found that we needed to allow for asymmetries in goal-congruent and incongruent evidence to capture our effects. Our winning model, the Goal-oriented Asymmetric Likelihood (GOAL) model, allowed *σ*_high_ and *σ*_low_ to vary independently. This meant that high and low evidence distributions, or target and non-target distributions as recognised in standard SDT, could have different variances (*σ*_Target_ ≠= *σ*_Nontarget_). Specifically, we found the goal-dependent asymmetry was *σ*_high_ *> σ*_low_ in the high evidence frame, and *σ*_high_ *< σ*_low_ in the low evidence frame. For the simulations in the Results section, we illustrated the model predictions by setting *σ*_high_ = 1.2 and *σ*_low_ = 1 within the high evidence frame, and *σ*_high_ = 1 and *σ*_low_ = 1.2 within the low evidence frame (2000 samples were extracted from the posterior distribution).

We further simulated the impact of various combinations of *σ*_High_ and *σ*_Low_ variances. A regression model was used to assess the impact of evidence on confidence (confidence ∼ ChosenEvidence + UnchosenEvidence). Both *σ*_high_ and *σ*_low_ were assigned values in the range [0.5, 1.9] with a step of 0.2. One thousand samples were extracted from each simulated model for each *σ*_high_ and *σ*_low_ combination. The contribution of chosen and unchosen evidence to confidence for various combinations of *σ*_high_ and *σ*_low_ in high evidence or low evidence frames is presented in Supplemental Figure S1.

Finally, we simulated a third model, the response-congruent heuristic model (HM;^10^) which was proposed to characterise evidence biases on confidence. In this model, confidence is generated by discarding the evidence supporting the unchosen option:

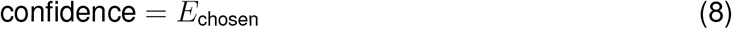

with *E*_chosen_ the evidence of the chosen option. For HM simulations, the model architecture is identical to EVM, except for the equation for confidence. All the model simulations were implemented in PyMC. Logistic and linear regressions were implemented using the sklearn and statsmodels toolboxes in Python. All values were normalized (z-scored) before fitting to the linear models.

### M.6 Model fitting

We used Markov chain Monte Carlo (MCMC) methods implemented in PyMC^64^ to sample posterior distributions of the parameters given the observed data. The actual evidence supporting the left and right options (*EV*_left_ and *EV*_right_, respectively), subjects’ confidence reports (*r*), choices (*c*) and reaction times were considered as observed variables in the models. The critical free parameters were the variances for the high and low evidence distributions. Priors were specified over these variances using normal distributions:

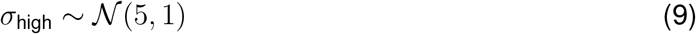

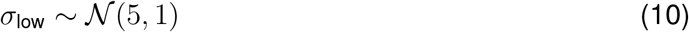

Although the normal distribution has support over negative values, the posterior distributions for *σ*_high_ and *σ*_low_ were consistently over zero across all chains and model posterior samples, thus confirming that this choice of prior did not pose a problem for estimation. As a robustness check, we additionally fitted all models using gamma-distributed priors over *σ*_high_ and *σ*_low_ (*α* = 6, *β* = 1.2), which are restricted to positive values. Results were consistent across both prior specifications (Figure S3), indicating that our findings are not an artifact of the choice of prior distribution.

We obtained the samples for the evidence observed for the left and right options (*E_l_*and *E_r_*, respectively) for the trial evidence for each one of the experiments:

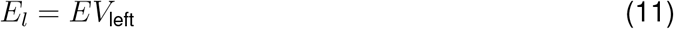

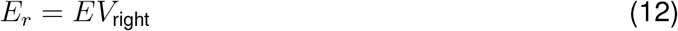

We defined a deterministic variable to estimate the decision variable (log-likelihood ratio from left and right options, *L*) as expressed in Equation 6.

To account for the stochasticity of human responses, we transformed the decision variable *L* with a logit function and estimated choices from a Bernoulli distribution.^34,52^ The parameter controlling the prior *β* was defined in terms of precision instead of standard deviation (precision *τ* = 0.001 → *σ* = 31.62). Therefore, choice was determined as:

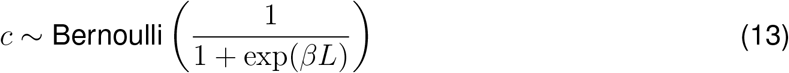

Where the prior for *β* was:

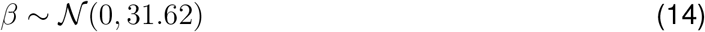

We additionally allowed modelled confidence to depend on the empirical reaction time (RT), which was included together with the estimated value of |*L*| (the magnitude of the log-likelihood term). The values of |*L*| and RT were entered into a logit function to generate a confidence rating on a 0–1 scale. The two additional weighting parameters (*β*_conf_ and *β*_RT_) were included in a deterministic expression to account for the contribution of both terms to the final reported confidence (overall confidence):

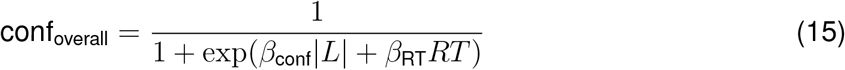

With priors:

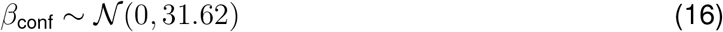

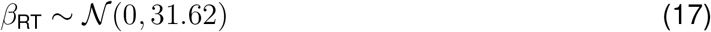

Following,^34^ the mapping between model confidence and the participants’ reported confidence allowed a small degree of imprecision (*σ* = 0.025) in subjects’ ratings, roughly equivalent to grouping continuous ratings made on a 0–1 scale into ten bins.

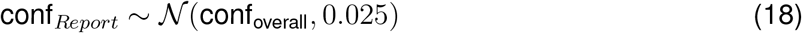

A schematic of this model is presented in Figure 6A. The fitting was performed at a pooled-participant level. *EV*_left_, *EV*_right_ were z-score normalised. Confidence was normalised at the participant level and transformed to the 0–1 range (*conf_Report_*).

In Experiment 1, the model used the willingness to pay reported by the participants for each food item to define the evidence means (*EV*_left_ and *EV*_right_). In Experiment 2, *EV*_left_ and *EV*_right_ corresponded to the number of dots presented in the left and right patches, respectively. In Experiment 3, we used the number of dots moving to the right or left to define evidence means. Finally, in Experiment 4, the number of auditory clicks presented in the left and right speakers defined the evidence means. Evidence values were normalised using z-scores at the participant level. Model structure and priors for the parameters were identical across experiments. While *σ*_high_ and *σ*_low_ were considered free parameters in the fitting process, *µ*_High_ and *µ*_Low_ were extracted from a median split on the participant’s evidence distribution. For the value-based decision Experiment 1, we used the distribution of subjective values reported by participants prior to the experiment. For experimenter-controlled perceptual stimuli in Experiments 2, 3 and 4, *µ*_high_ and *µ*_Low_ were extracted from dot number, coherence levels, and auditory click number distributions presented to participants. The distribution of evidence and median splits are presented in Supplemental Figure S2. The average of the values higher than the median was assigned as *µ*_High_ and the average of evidence values under the median as *µ*_Low_, and used as constants in the models.

The model was fitted to even-numbered trials. We fitted each model by generating 4,000 samples in each of the 4 chains using the NUTS sampling method.^65^ We discarded 1000 for tuning of the sampler. Convergence was assessed using the Gelman–Rubin statistic (|*R*^^^− 1| *<* 0.05), showing good convergence for the parameters of interest *σ*_high_ and *σ*_low_. Regression analyses for simulated trials and test trials were performed using the statsmodels package in Python. For choice models, we predicted the log odds ratio of selecting the item appearing on the right. As in the behavior analysis, confidence model included RT, *confidence_simulated_* ∼ *RT* + *ChosenEv* + *UnchosenEv*. The number of posterior samples for the analysis depended on the number of trials used to fit the model: Experiment 1: 93,000 total samples (1,860 trials × 50); Experiment 2: 96,000 total samples (1,920 trials × 50); Experiment 3: 162,900 total samples (3,258 trials × 50); Experiment 4: 72,000 total samples (1,440 trials × 50).

### M.7 Data analysis: pupillometry experiment

#### M.7.1 Pupil Dilation Pre-processing

Eye-tracking data were processed using DataViewer 4.2.1 (SR Research), generating reports containing time series of pupil area measurements for all participants. The data were subsampled to 100 Hz. Pre-processing included interpolation of eye blinks^25,28^ and removal of blink– and saccade-related artefacts through deconvolution followed by linear regression.^66^

Each trial’s pupil trace was z-scored and baseline-corrected by subtracting the mean pupil area from the 2 seconds preceding the onset of the auditory stimulus. For visualisation, pupil traces were time-locked to key events—stimulus onset and response time—and epoched accordingly. The pre-processed data were then entered into a general linear model (GLM) analysis.

#### M.7.2 GLM Analysis

To examine how pupil dynamics reflected decision evidence, we fitted a General Linear Model (GLM) at each time point of the pupil time series. Analyses were focused around the moment of choice, using a 4 s window (2 s before and 2 s after the response). Trials were analysed separately for the *High Clicks* and *Low Clicks* frames.

For each participant, we extracted the pupil trace aligned to response time and fitted a GLM predicting relative pupil size from the following regressors:

- Chosen evidence (number of clicks on the selected side),
- Unchosen evidence (number of clicks on the non-selected side),
- Reaction time (RT), and
- Gaze position (*X*–*Y* coordinates at each time point).

The inclusion of gaze position controlled for potential nuisance effects.^28^

At the group level, we focused on the parameter estimates describing the effect of chosen evidence on pupil size (Figure 7). Statistical significance was assessed through a permutation analysis: for each participant (*n* = 32) and time sample (400 total), we fitted 50 GLMs using permuted datasets to generate a null distribution of group-level parameters for both frames. The difference in chosen-evidence parameters between the High and Low Clicks frames was computed to obtain a null distribution for the contrast.

Significance of the observed difference was determined by comparing the empirical parameter contrast to this null distribution. P-values were corrected for multiple comparisons using a false discovery rate (FDR) threshold of *α* = 0.01. Only clusters of at least six contiguous significant time samples were considered reliable.^15^

## Data Availability

All data and code used for this study will be made available upon acceptance of the manuscript.

## Author Contributions

PS and BDM conceived the project and developed the experiments. PS and MZ conducted the experiments. PS performed data analyses and model fitting, with assistance and input from SF and BDM. PS, SF and BDM discussed the results and wrote the manuscript, with input from MZ.

## Acknowledgments

We thank Matan Mazor, Yumeya Yamamori, Giuseppe Castegnetti, Amy Benson and Mihaela Nemes for valuable discussions and input into this project. PS was funded by the Chilean National Agency for Research and Development (Graduate student scholarship – DOCTORADO BECAS CHILE/2017 – 72180193).

## Supplementary Figures

**Figure S1:**
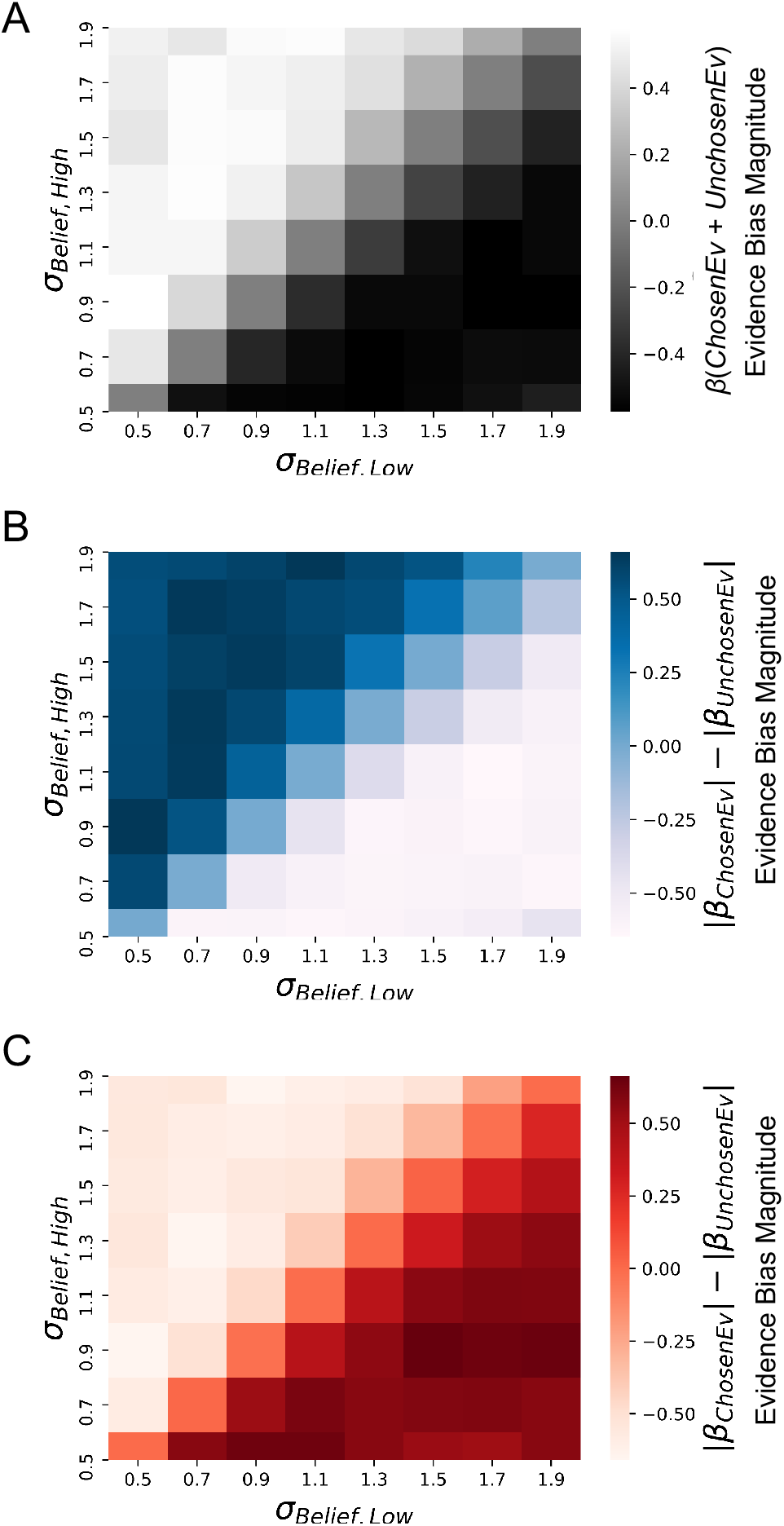
Representation of evidence bias effects for multiple combinations of belief variance. For each combination of belief variance (*σ*_Low_, *σ*_High_), 1000 trials were simulated. These simulations were used to fit linear regression models predicting confidence. (A) Effect of ΣEvidence on confidence shows how simulations using *σ*_Low_ *< σ*_High_ generated a positive effect of ΣEvidence on confidence, while *σ*_Low_ *> σ*_High_ produced a negative effect of ΣEvidence. (B–C) Similar simulations, but now presenting the imbalance in the contribution of chosen and unchosen evidence to confidence. In simulations where the higher evidence option was selected (B), a bigger contribution of chosen evidence to confidence is observed when *σ*_Low_ *< σ*_High_. (C) On the other hand, when the lower evidence option was selected, simulations with *σ*_Low_ *> σ*_High_ generated an over-weighting of the chosen option in the confidence signal. Importantly, note that in all panels, equal-variance simulations predicted no effect of ΣEvidence or imbalance of chosen and unchosen evidence.

**Figure S2:**
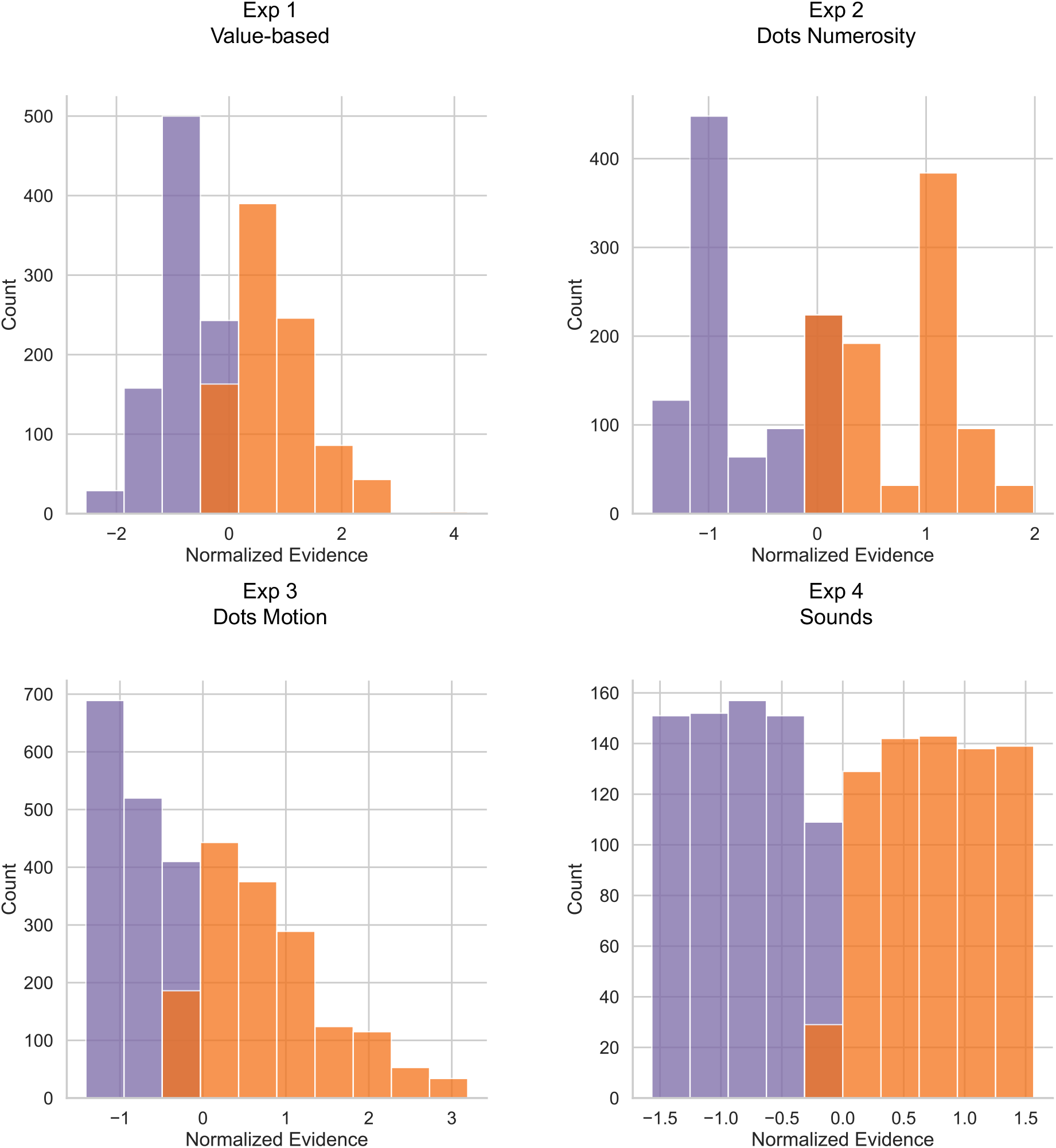
Evidence distribution observed in the experiments. The median values of the distributions was selected to separate high from low distributions. The average values *µ_High_* and *µ_Low_* were calculated from these empirical distributions and used in the model.

**Figure S3:**
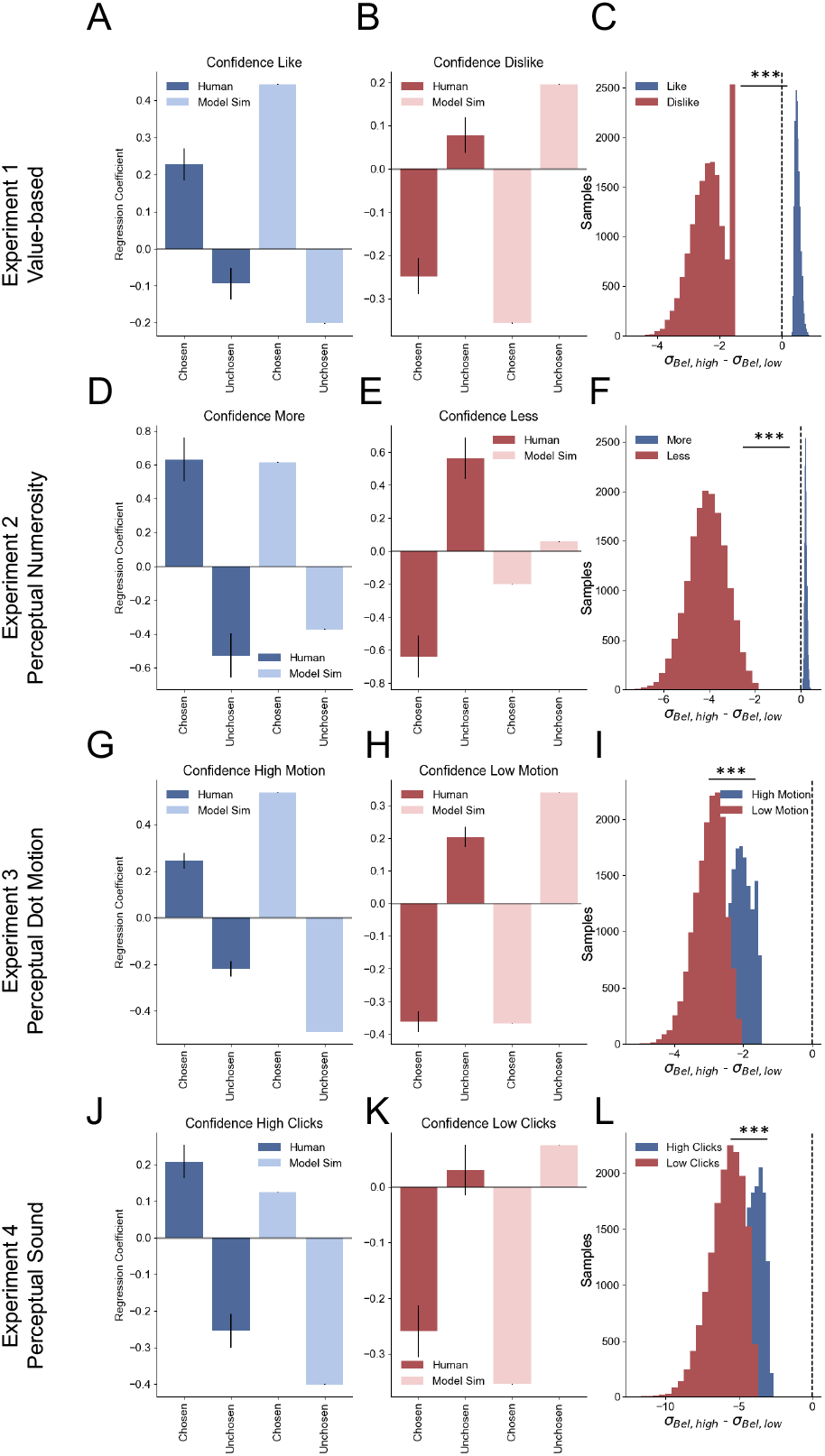
GOAL model fit to decision making experiments. *Results using gamma distribution as priors for σ_bel_*. Simulations replicate the pattern of results obtained from the regression analysis on participants’ behaviour and the GOAL model simulations using Normal priors for *σ*_bel_. (A–C) Experiment 1: model simulations for like (A) and dislike (B) frames. Specifically, the model captures the frame-dependent variation of the chosen and unchosen evidence effect on confidence. Latent parameters, *σ*_high_ and *σ*_low_ are distinct depending on the frame, with the variance of the goal-relevant alternative relatively higher (C). (D–F) Experiment 2: model simulations for most (D) and fewest (E) frames, replicate the behavioural effect of chosen and unchosen on confidence, and its interaction with the frame. Belief variance also presents the expected asymmetry, with higher variance for the goal-relevant option (F). (G–I) Experiment 3: model simulations for high (G) and low dot motion frames (H). Simulations replicate human behaviour in confidence. The values of *σ*_bel_ for high and low distributions were significantly different between frames (I). (J–L) Experiment 4: The model simulations captured the pattern of confidence effects for the auditory experiment in the high (J) and low (K) clicks frames. The variance distributions *σ*_bel_ for high and low clicks were also significantly different depending on the frame. Linear regression models are presented for confidence in participants and model simulations (pooled linear regression). *σ*_bel,high_ – *σ*_bel,low_ distributions in panels were generated from sampling posterior distributions of the fitted models.

